# Summarizing RNA Structural Ensembles via Maximum Agreement Secondary Structures

**DOI:** 10.64898/2026.02.25.708069

**Authors:** Xinyu Gu, Stefan Ivanovic, Daniel W. Feng, Mohammed El-Kebir

## Abstract

Summarizing a collection **P** of related RNA secondary structures is a key challenge in applications like evolutionary analysis, alternative fold studies and mRNA vaccine design. This requires both clustering the input structures into similar groups and identifying the core structural motifs on which they agree or differ. Existing methods fail by focusing on only one of these goals: clustering methods do not output shared motifs, while consensus methods overlook the structural diversity present in the collection. Here, we introduce the Maximum Agreement Secondary Structures (MASS) problem, which seeks the largest set *F* of structural features present in **P** that partition the input structures into a user-specified number *τ* of distinct clusters. We prove that MASS is NP-hard and also establish its equivalence to a constrained binary matrix projection problem. We present an exact integer linear program, an exact combinatorial algorithm, and a scalable beam-search heuristic. Using simulations we demonstrate the performance of these exact algorithms and heuristics relative to baseline methods that focus on either clustering or identifying a single consensus tree. On real data, we demonstrate that MASS identifies conserved scaffolds in conformational datasets, reveals conserved structural motifs in different species within RNA families, and recovers shared structural features among synonymous transcripts encoding the same protein. MASS provides a general and interpretable framework for summarizing RNA structural organization.

## 1 Introduction

RNA secondary structure plays a major role in determining RNA function, affecting transcriptional attenuation, translation, RNA stability, localization, and molecular interactions [1]. These regulatory effects are frequently encoded in discrete structural motifs, which can function independently of the underlying nucleotide sequence [1, 2, 3]. There are three key applications where one is given a collection **P** of aligned RNA structures *P*_1_, …, *P*_*m*_ and seeks to identify shared structural motifs. First, we could be given a collection **P** of alternative folds of the same RNA sequence **v** [4]. Second, **P** may correspond to an RNA family composed of different evolutionary-related species [5]. Third, for mRNA vaccine design for a given target protein sequence **w**, we could be given a collection (**P, V**) of RNA structure-sequence pairs that encode for the same target protein **w** [6, 7, **8]**. The key underlying challenge for these applications is developing a method to summarize the collection **P**, simultaneously identifying a clustering of the *m* structures into groups of similar structures as well as identifying the core structural motifs on which its members agree and differ.

Several methods have been proposed to address this challenge, focusing either on clustering the structures or on identifying a single consensus structure. The first category of methods have been applied to cluster RNA sequences and structures across multiple RNA families and to recover meaningful functional groupings [9], and typically proceed in a two-step fashion. First, these methods obtain an *m* × *m* distance matrix *D* by pairwise comparison of the aligned structures using the symmetric difference of base pairings [10] or by pairwise sequence-structure alignment using algorithms such as bpRNA-align [9], RNAforester [11] and BEAGLE [12]. Then, the distance matrix *D* is used for agglomerative clustering, after which the dendogram is cut to obtain the final clustering. The downside of this first category of methods is that they do not output the precise set of structural motifs that best summarize the input structures **P**. The second class of methods is based on identifying a consensus RNA structure that best represents the input structures. One recent method does so by representing the input structures as trees, thereby disallowing pseudoknots, and solving a median problem [13]. Specifically, this method seeks to identify a tree/structure *P* ^∗^ that minimizes the sum of distances to each input tree/structure *P*_1_, …, *P*_*m*_ where the distance is a tree distance measure such as Robinson-Foulds [14]. The downside of this second class of approaches is that a single consensus structure might be an inadequate representation of the full structural diversity of the input structures **P** — this is especially problematic when imposing additional constraints on the consensus *P* ^∗^ such as being pseudoknot free. Alternative consensus approaches, such as Maximum Expected Accuracy (MEA) estimation [15, 16], are specific to thermodynamic ensembles of a single sequence (application 1), while covariance-based methods like RNAalifold [17], PETfold [18] and CentroidAlifold [19] are restricted to homologous sequence alignments (application 2).

We introduce the Maximum Agreement Secondary Structures (MASS) problem, which takes as input a collection **P** of RNA secondary structures and identifies the largest set of structural features that can be jointly supported across the input, while allowing the structures to partition into a limited number of distinct structural states (Fig. 1). MASS assumes an alignment of the input, but there is no assumption about the inputs being pseudoknot free. MASS summarizes the shared structural core and explicitly identifies variable structural elements. In this work, we formalize the MASS problem, show that it is equivalent to a constrained matrix projection problem and establish NP-hardness. We present two exact algorithms, one based on integer linear programming and the other a combinatorial algorithm. In addition, we introduce a beam-search heuristic that scales to large instances. We demonstrate that MASS identifies conserved scaffolds in conformational datasets, reveals conserved structural motifs in different species within RNA families, and recovers shared structural features among synonymous transcripts encoding the same protein. MASS provides a general and interpretable framework for summarizing RNA structural organization.

**Figure 1:**
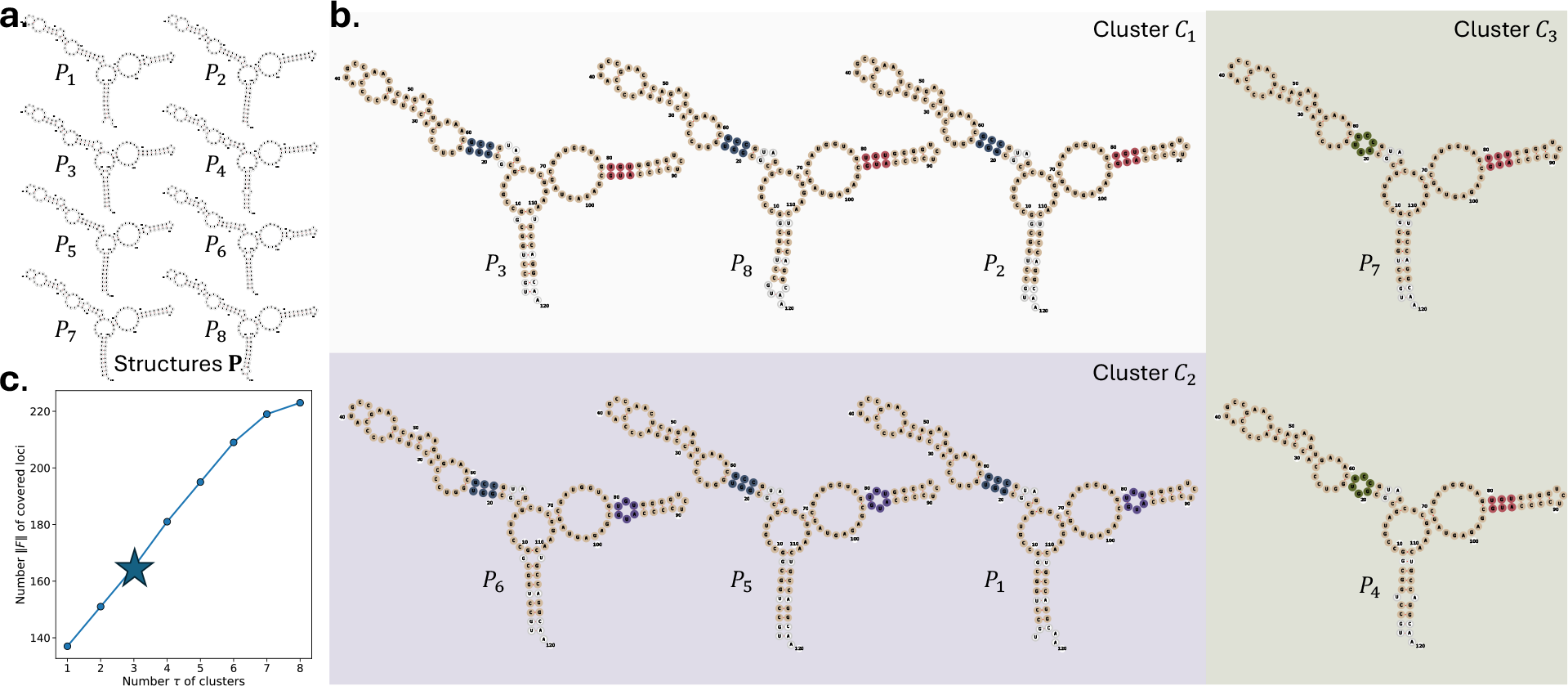
Overview of Maximum Agreement Secondary Structures. (a) Given a collection **P** of *m* RNA structures and number *τ* [*m*] of clusters, we seek to select a subset *F* of features with maximum number ∥*F* ∥ of covered loci that induces at most *τ* clusters. (b) Solution with *τ* = 3 clusters; with nucleotide coloring indicating cluster membership (yellow: present in all clusters; blue: *C*_1_, *C*_2_; red: *C*_1_, *C*_3_, purple: *C*_2_; green: *C*_3_). (c) There is a tradeoff between the number ∥*F* ∥ of covered loci and the number *τ* of clusters.

## 2 Problem Statement

We are given *m* align RNA secondary structures **P** = (*P*_1_, …, *P*_*m*_). Secondary structures can be represented in one of two ways. First, in the *base pairing (BP) representation*, one looks at the set of complementary base pairings present in each structure (Fig. S1). Second, in the *secondary structure element (SSE) representation*, one extracts all secondary structure elements, which include stacking, hairpin, bulge loops, interior loops and multiloops, recording the precise set of nucleotide loci comprising each element (Fig. S1). We define the two representations as follows.

### Definition 1.

*The* base pairing (BP) representation *of a secondary structure P with ℓ*_bp_(*P*) *base pairings of an RNA sequence with n nucleotides equals* 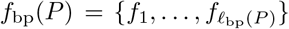 *where each f* ∈ *f*_bp_(*P*) *corresponds to a unique base pairing* (*p, q*) ∈ [*r*]*×*[*r*] *(where* [*r*] = {1, …, *r*}). *The number* ∥*f* ∥ *of covered loci equals* 2 *for each f* ∈ *f*_bp_(*P*).

### Definition 2.

*The* secondary structure element (SSE) representation *of a secondary structure P with ℓ*_sse_(*P*) *secondary structure elements of an RNA sequence with n nucleotides equals* 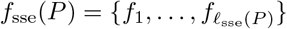 *where each secondary structure element f* ∈ *f*_sse_(*P*) *corresponds to a pair* (*X, Y*) *of sets where X* ⊆ [*r*] *×* [*r*] *indicates the base pairings present in f and Y* ⊆ [*r*] *is the set of unpaired nucleotides that are part of f* . *The number* ∥*f* ∥ *of covered loci equals* · |*X*| + |*Y* | *for each secondary structure element f* = (*X, Y*).

In the following, we will omit the precise type of representation and given a representation *f* (*P*) = *f*_1_, …, *f*_*ℓ*_ simply refer to each comprising element *f*_*j*_ as a feature. Our goal is to summarize the collection **P** by maximizing feature agreement. To that end, we introduce the following definition.

### Definition 3.

*Given structures* **P** = (*P*_1_, …, *P*_*m*_), *the secondary structure feature space* ℱ (**P**) *equals* 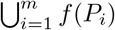.

Generalizing the previous notation, we define ∥*F* ∥ = ∑_*f*∈*F*_ ∥*f* ∥ to denote the total number of loci present in *F* ⊆ ℱ (**P**). Note that the same locus might occur in multiple features in *F*, and be counted multiple times. The key observation is that each subset *F* ⊆ ℱ (**P**) of features induces a unique partition of the *m* structures, as defined next.

### Definition 4.

*Given structures* **P** = (*P*_1_, …, *P*_*m*_) *and features F* ⊆ ℱ (**P**), *the F* -induced partition *C*(**P**, *F*) *is a partition of* [*m*] = {1, …, *m*} *such that for all clusters C* ∈ *C*(**P**, *F*) *and all comprising structures P*_*i*_, *P*_*j*_ ∈ *C it holds that P*_*i*_ *and P*_*j*_ *agree on features F, i*.*e. f* (*P*_*i*_) ∩ *F* = *f* (*P*_*j*_) ∩ *F* .

The goal is to select a subset *F* ⊆ ℱ (**P**) of features that ideally optimize two criteria. First, we wish to maximize the number ∥ *F* ∥ of covered loci. Second, we wish to minimize the number | *C*(**P**, *F*) | of resulting clusters. One can maximize the first objective, the number of covered loci, by simply setting *F* = ℱ (**P**) but this would come at the expense of the second objective, yielding a clustering *C*(**P**, *F*) of maximum size (equal to *m* provided the *m* input RNA structures are distinct). On other hand, minimizing the second objective, the size of the clustering is trivial, choosing any individual feature *f* ∈ ℱ and setting *F* = {*f* } would yield a clustering *C*(**P**, *F*) of size at most 2 (equal to 1 if all input structures contain *f*). We choose to model this tradeoff via a budget constraint with parameter *τ* ∈ ℕ specifying the maximum allowed number of clusters, i.e. |*C*(**P**, *F*)| ≤ *τ* for any choice of *F* ⊆ ℱ (**P**).

### Problem 1.

(Maximum Agreement Secondary Structures (MASS)). *Given m align RNA structures* **P** *and parameter τ* ∈ [*m*], *select a subset F* ⊆ ℱ (**P**) *of features inducing a partition C*(**P**, *F*) *of at most τ clusters such that the number* ∥*F* ∥ *of covered loci is maximized*.

## 3 Combinatorial Characterization and Complexity

Due to space constraints, proofs are relegated to Section A.

### 3.1 Combinatorial Characterization

We begin by showing that Problem 1 is equivalent to a binary matrix projection problem. Specifically, for a given *m × n* binary matrix *B* we denote with *B*[*N*] the binary matrix obtained by (i) retaining columns *N* and removing the remaining [*n*] \ *N*, followed by (ii) removing duplicate rows.

#### Problem 2.

(Binary Matrix Column Selection and Projection (BMCSP)). *Given a binary matrix B* ∈ { 0, 1 } ^*m×n*^ *and parameter τ* ∈ [*m*], *find the largest subset N* ⊆ [*n*] *of columns that results in a matrix B*[*N*] *with at most τ rows*.

We have the following important theorem.

#### Theorem 1.

BMCSP *is equivalent to* MASS.

We show this by first defining how to reduce a MASS instance (**P**, *τ*) to a corresponding BMCSP instance (*B, τ*). We note that the reduction works for both the BP and SSE representation of secondary structures as the choice of representation only alters the feature space ℱ (**P**).

#### Definition 5.

*Let* (**P**, *τ*) *be a* MASS *instance composed of m structures* **P** = (*P*_1_, …, *P*_*m*_) *and an integer τ* ∈ ℕ. *The corresponding* BMCSP *instance* (*B, τ*) *consists of a binary matrix B with m rows and* ∥ ℱ (**P**) ∥ *columns, by introducing, for each feature F* ∈ ℱ (**P**), ∥*F* ∥ *identical columns to B with a* 1 *in row i* ∈ [*m*] *if F is present in structure P*_*i*_ *and a* 0 *otherwise*.

We refer to Fig. S2 for an example reduction. We now prove the correctness of the reduction.

#### Proposition 1.

*Let B* ∈ { 0, 1 } ^*m×n*^ *be a binary matrix obtained from secondary structures* **P** *following Definition 5. Then, there exists an optimal* BMCSP *solution N composed of k columns such that B*[*N*] *has τ rows if and only if there exists an optimal* MASS *solution F* ⊆ ℱ (**P**) *covering* ∥ *F* ∥ = *k loci and inducing a partition C*(*F*, **P**) *of τ clusters*.

We now define and prove the correctness of the reverse reduction from a BMCSP instance (*B, τ*) to a corresponding MASS instance (**P**, *τ*).

#### Definition 6.

*Let* (*B, τ*) *be a* BMCSP *instance composed of a binary matrix B* ∈ {0, 1}^*m×r*^ *and an integer τ* ∈ ℕ. *The corresponding* MASS-BP *instance* (**P**, *τ*) *consists of m structures where each structure P*_*i*_ *(where i* ∈ [*m*]*) has length* 2*r and consists solely of base pairings of the form* (*j*, 2*r* − *j* + 1) *if matrix entry b*_*i,j*_ = 1 *(where j* ∈ [*r*]).

#### Proposition 2.

*Let* **P** *be the* MASS-BP *instance obtained from matrix B (without all* 0*s columns) following Definition 6. Then, there exists an optimal MASS-BP solution F* ⊆ ℱ_bp_(**P**) *covering* ∥*F* ∥ = 2*k loci and inducing* | *C*(*F*, **P**) | = *τ clusters if and only if there exists an optimal* BMCSP *solution N such that B*[*N*] *has k columns and τ rows*.

### 3.2 Complexity

We have the following hardness results.

#### Theorem 2.

MASS *is NP-hard*.

#### Theorem 3.

BMCSP *is NP-hard*.

We prove this by giving a polynomial-time reduction from Independent Set to Binary Matrix Column Selection and Projection (BMCSP), which, in turn, reduces to Maximum Agreement Secondary Structures (MASS) in polynomial time (as previously shown in Proposition 2). We begin by recalling the independent set problem.

#### Problem 3.

(Independent Set). *Given an undirected, connected graph G* = (*V, E*) *and an integer k* ∈ ℕ, *determine whether there exists a subset I* ⊆ *V of size k such that no two vertices in I are adjacent*.

This problem remains NP-complete when assuming the graph is connected [20].

#### Definition 7.

*Let* (*G* = (*V, E*), *k*) *be an instance of* Independent Set *where V* = {*v*_1_, …, *v*_*n*_} *and E* = {*e*_1_, …, *e*_*m*_}. *The corresponding* BMCSP *instance* (*B, τ*) *consists of a binary matrix B* ∈ {0, 1}^3*m×n*^ *such that (i) each vertex v*_*j*_ ∈ *V corresponds to column j of matrix B; (ii) each edge e*_*i*_ = (*v*_*j*_, *v*_*j*_*′*) ∈ *E corresponds to the three rows* {3(*i* − 1) + 1, 3(*i* − 1) + 2, 3(*i* − 1) + 3} *inducing a submatrix of B corresponding composed of entries* (1, 0), (0, 1), (1, 1) *at columns* {*j, j*^*′*^} *and having* 0 *values at the remaining columns; and (iii) τ* = *k* + 1.

This construction, illustrated in Fig. S2, takes polynomial time. Finally, we have the following lemma proving correctness of the reduction.

#### Lemma 1.

*Let* (*B, τ*) *be an instance of* BMCSP *obtained from a* Independent Set *instance* (*G, k*) *following Definition 7. Then, there exists an independent set I of G composed of* |*I*| = *k vertices if and only if there exists a subset N of columns inducing a submatrix B*[*N*] *composed of* |*N* | = *k columns and at most τ* = *k* + 1 *rows*.

## 4 Methods

Leveraging the equivalence and polynomial-time reduction from MASS to BMCSP (see Proposition 1), let *B* = [*b*_*i,j*_] be the *m* × *n* binary matrix obtained from RNA structures **P** = (*P*_1_, …, *P*_*m*_) following the reduction given in Definition 5. The goal is to find the largest subset *N* ⊆ [*n*] of columns that induces a matrix *B*[*N*] with at most *τ* rows. We do so using an integer linear program (ILP) in Section 4.1, followed by an exact, combinatorial algorithm in Section 4.2, which we adapt into a beam-search heuristic in Section 4.3. Finally, we discuss two baseline methods in Section 4.4. We implemented these methods in Python and used Gurobi version 12.0.3 for the ILP. Our software is open source and available at https://github.com/elkebir-group/MASS.

### 4.1 Integer Linear Program

To obtain an ILP, we must model (i) column selection corresponding to *N*, (ii) enforce that *B*[*N*] consists of at most *τ* rows and (iii) additional optimizations. This leads to the following ILP with *O*(*n*+*m*^2^) variables, including *O*(*n*+*m*^2^) binary variables, and *O*(*n* + *m*^2^) constraints.

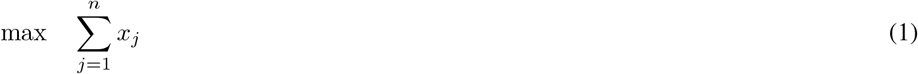

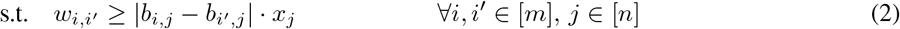

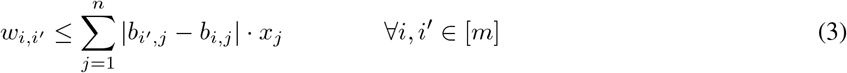

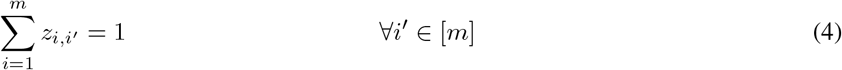

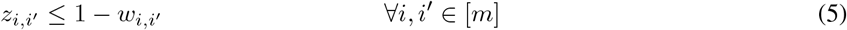

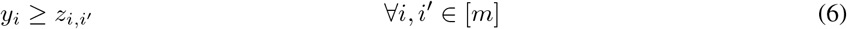

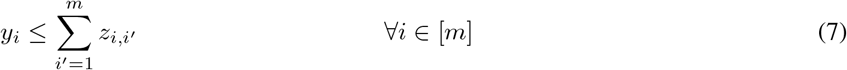

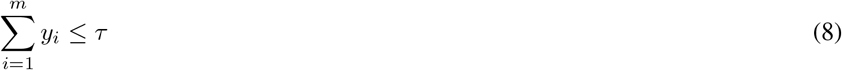

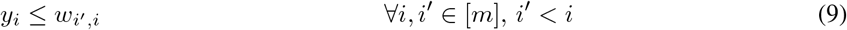

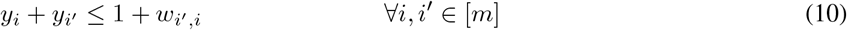

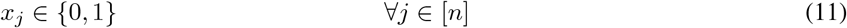

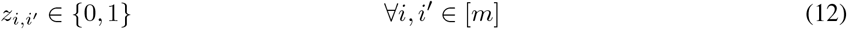

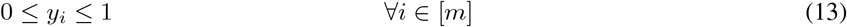

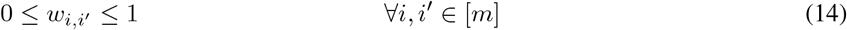

In the following, we explain the variables and constraints of the ILP.

#### (i) Column selection *N*

We introduce binary variables **x** ∈ {0, 1}^*n*^ such that *x*_*j*_ = 1 if and only if column *j* ∈ *N* . We express the goal to maximize |*N* | in (1).

#### (ii) Clustering of rows

A key constraint of BMCSP (Problem 2) is that *B*[*N*] must consist of at most *τ* rows. To model this using linear constraints, we begin by introducing binary variables **w** ∈ { 0, 1 } ^*m×m*^ such that *w*_*i,i*_*′* = 1 if and only if there exists a column *j* ∈ [*N*] such that *b*_*i,j*_ ≠ *b*_*i*_*′,j* . This enforced by (2) and (3). We note that these two constraints are linear as the absolute difference | *b*_*i*_*′,j* −*b*_*i,j*_ | w.r.t. two constants can be precomputed.

Next, we introduce the concept of a “representative row” such that each set of identical rows w.r.t. columns *N* is represented by exactly one member of that set. We introduce binary variables **y** ∈ { 0, 1 } ^*m*^ such that *y*_*i*_ = 1 if and only if row *i* is a representative row as well as binary variables **z** ∈ { 0, 1 ^*m×m*^ } such that *z*_*i,i*_*′* = 1 if and only if row *i* is a representative of row *i*^*′*^. We enforce this using several constraints. First, in (4), we have that each row *i*^*′*^ must be represented by exactly one row (can be row *i*^*′*^ itself). Second, in (5), we have that a row *i* cannot represent *i*^*′*^ if the two rows differ w.r.t. a column in *N* (i.e., *w*_*i,i*_*′* = 1). Third, in (6), we have that a row *i* can only represent *i*^*′*^ if *i* is a representative row itself. Fourth, in (7), we have that a row *i* cannot be a representative if there are rows *i*^*′*^ that it represents. Finally, (8) corresponds to the key constraint that there can only be at most *τ* representative rows.

#### (iii) Optimizations

We note that constraints discussed thus far ensure correctness. However, there is still room for improvement. To further strengthen the formulation, we include several optimizations. First, we break symmetry by requiring that only the first appearing row is the representative of a set of rows that are identical w.r.t. columns *N* . This is modeled by (9). Second, it cannot be that two identical rows are two distinct representatives, as modeled by (10). Finally, we note that integrality of **w** is implied by integrality of **x** [see (11)], and that integrality of **y** is implied by integrality of **z** [see (12)]. Thus, it suffices to require that these two sets of variables are continuous, as in (13) and (14).

### 4.2 Exact Combinatorial Algorithm

Given the input matrix *B* ∈ { 0, 1 } ^*m×n*^, we denote with *N* ⊆ [*n*] a subset of columns of *B*. We denote with ℳ a partition of all rows [*m*] of *B*. Akin to Definition 4, each subset *N* of columns ℳ [*n*] induces a partition (*N*) of the *m* rows as follows.

#### Definition 8.

*Given a matrix B* ∈ {0, 1}^*m×n*^ *and a subset N of columns* [*n*], *the N* -induced row partition ℳ (*N*) *consists of subsets M*_*i*_ ∈ ℳ (*N*) *of rows with either all* 0 *or all* 1 *entries at each column of N, i*.*e*. 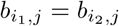 *for all rows i*_1_, *i*_2_ ∈ *M*_*i*_ *and columns j* ∈ *N* .

This can be computed in *O*(*m* |*N* |) time by first extracting a bit string **x**_*i*_ ∈ { 0, 1 } ^|*N*|^ of each row *i* ∈ [*m*] composed of the entries at columns *N* . These bit strings **x**_1_, …, **x**_*m*_ can then be hashed or sorted using radix sort to extract the partition ℳ (*N*). Next, we show that each partition ℳ of [*m*] induces the largest subset *N* (ℳ) of columns as follows.

#### Definition 9.

*Given a matrix B* ∈ { 0, 1} ^*m*×*n*^, *a partition* ℳ *of*, [*m*] *the* ℳ-induced column set *N* (ℳ) *equals the intersection* 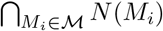 *where N* (*M*_*i*_) *is the set of all columns composed of either all* 0 *or all* 1 *entries at rows/part M*_*i*_ *of* ℳ, *i*.*e. N* (*M*_*i*_) *is composed of all columns j* ∈ [*n*] *such that* 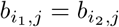 *for all rows i*_1_, *i*_2_ ∈ *M*_*i*_.

We note that *N* (ℳ) = ∅ if there is no subset *N* ^*′*^ of columns [*n*] that induces a partition ℳ (*N* ^*′*^) of which ℳ is a refinement. Given matrix *B* and a partition ℳ, we can compute the ℳ *-induced column set N* (ℳ) in *O*(*mn*) time by simply iterating over all columns *j* ∈ [*n*] and then checking whether each *M*_*i*_ ∈ ℳ is constant in column *j* (i.e., not consisting of both a 0 and 1 entry). We now prove that *N* (ℳ) is indeed the largest set of columns that induces ℳ.

#### Proposition 3.

*Let N* ^∗^ = *N* (ℳ) *be the* ℳ *-induced column set of a given partition* ℳ *of rows* [*m*] *of matrix B* ∈ {0, 1}^*m×n*^. *Then, for every subset N* ^*′*^ ⊆ [*n*] *of columns such that* ℳ (*N* ^*′*^) *equals* ℳ *it holds that N* ^*′*^ ⊆ *N* ^∗^.

As optimal solutions to BMCSP must maximize |*N* |, the above proposition implies a simple exhaustive search algorithm. That is, we can simply enumerate all partitions ℳ = {*M*_1_, …, *M*_*ℓ*_} consisting of *ℓ*^*′*^ ≤ *τ* non-empty parts and generate the corresponding ℳ -induced column set *N* (ℳ), and simply return the partition ℳ |*N* (ℳ)|. The number of partitions of [*m*] of size *ℓ* equals the Stirling number *S*(*m, ℓ*) of the second kind, which, for constant *ℓ*, equals *O*(*ℓ*^*m*^) asymptotically. Reconstructing the corresponding column set *N* (ℳ) for each enumerated that maximizes partition takes *O*(*mn*) time. Therefore, we have a total running time of 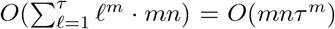, rendering this algorithm very impractical. We provide the pseudocode in Algorithm 1. To obtain a more efficient algorithm, we prove the following theorem.

#### Theorem 4.

*Let* ℳ = {*M*_1_, …, *M*_*τ*_ } *be a partition of rows* [*m*] *of matrix B* ∈ {0, 1}^*m×n*^ *such that there exists a subset N* ^*′′*^ ⊆ [*n*] *of columns where* ℳ (*N* ^*′′*^) = ℳ. *Then, there exists a subset N* ^*′*^ ⊆ [*n*] *of columns of size* |*N* ^*′*^| ≤ *τ* −1 *such that N* ^*′*^ *also induces partition* ℳ, *i*.*e*. ℳ (*N* ^*′*^) = ℳ.

After establishing Theorem 4, which shows that any partition ℳ with | ℳ| = *τ* parts can be obtained from *τ* − 1 columns, a naive algorithm for BMCSP follows. Given the threshold *τ* ∈ ℕ, we enumerate all subsets *N* ⊆ [*n*] such that | *N* | ≤ *τ* − 1. For each subset *N* of size *ℓ* = | *N* |, we compute the *N* -induced row partition ℳ (*N*); this takes *O*(*mℓ*) time per subset. If | ℳ (*N*) | ≤ *τ* then we score *N* by the number | *N* ℳ ((*N*)) | of columns of the largest subset *N* (ℳ (*N*)) of columns that also induces row partition ℳ (*N*). Each score evaluation takes *O*(*mn*) time. Finally, the algorithm outputs the subset achieving the maximum score. Assuming *τ* is constant, the algorithm takes 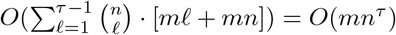 time. We provide the pseudocode in Algorithm 2.

As an optimization, observe that two different subsets *N, N* ^*′*^ of columns can, and often do, produce the same induced partition of rows, i.e. ℳ (*N*) = ℳ (*N* ^*′*^). The algorithm that we discussed in the previous paragraph wastes effort by treating *N* and *N* ^*′*^ as distinct, even though from the perspective of row partitions they represent the same state. Thus, what matters is not which subset of columns was used, but which partition of the rows it induces. As such, we introduce the Max-Subset *τ* -Partitioning (MSTP) algorithm, which avoids redundant computations on equivalent column sets by instead iteratively constructing and deduplicating the partitions themselves. We start by initializing the list ℒ_current_ of partitions with just a single partition composed of a single part [*m*] containing all rows. Then, starting with *ℓ* = 1, each iteration *ℓ* ≤ *τ* − 1 refines the unique partitions from the previous step with every column *j* ∈ [*n*], discards any resulting partition larger than *τ*, and then sorts (via radix sort) and deduplicates the resulting list ℒ_next_ of *O*(*n*^*ℓ*^) candidates. This process takes *O*(*mn*^*ℓ*^) time per iteration, leading to a total of *O*(*mn*^*τ*−1^) for all *τ* − 1 iterations. At the end of the final iteration *ℓ* = *τ* − 1, we score the final list ℒ_current_ of at 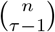 most unique partitions by computing the maximum number | *N* (ℳ) | of induced columns among all enumerated partitions ℳ. We return the partition ℳ^∗^ and corresponding largest column set *N* (ℳ^∗^). This final evaluation takes *O*(*mn* · *n*^*τ*−1^) = *O*(*mn*^*τ*^) time and thus dominates the overall running time. We provide pseudocode in Algorithm 3. While this algorithm has the same worst-case asymptotic running time as Algorithm 2, we expect it to perform better in practice. Moreover, this algorithm is amenable to beam search, as we will discuss next.

### 4.3 Beam-search Heuristic

To obtain a beam-search heuristic, we introduce a parameter *w* ∈ ℕ that controls the beam width, i.e. the number of top-*w* scoring partitions retained after deduplication. Specifically, we maintain a list ℒ_current_ of score-partition pairs by adapting the MSTP (Algorithm 3) to compute the score *N* (ℳ^*′*^) of ℳ^*′*^-induced column set for each newly constructed partition ℳ^*′*^. For each iteration *ℓ* ∈ { 1, …, *τ* − 1 }, which starts with | ℒ_current_ | ≤ *w* partitions, this refine- and-score step takes *O*(*w*· *n* ·*mn*) = *O*(*wmn*^2^) time. Given that the resulting list | ℒ_next_ | contains *O*(*wn*) partitions, the sort-and-deduplicate step takes *O*(*wn* log(*wn*)) time. The total cost per iteration is the sum of these components, *O*(*wmn*^2^ + *wn* log(*wn*)). The *τ* − 1 iterations thus take a total of *O*(*τ* (*wmn*^2^ + *wn* log(*wn*))) time. In practice, we have *w* = *O*(*n*^*τ*−1^) as the number of unique partitions generated in *τ* − 1 iterations is bounded by *O*(*n*^*τ*−1^) and thus setting *w* to a larger value has no effect. When additionally assuming *τ* = *O*(1), which is the case in practice, the running time simplifies to *O*(*τwmn*^2^). We provide the pseudocode of this beam search procedure in Algorithm 4.

### 4.4 Baseline Methods

We consider two baseline methods. While these methods do not directly solve MASS (Problem 1), their output can be transformed into MASS output as we discuss in the following.

First, we use the results obtained from the median method introduced by Marchand *et al*. [13], which we refer to as `RNAconsensus`. Briefly, this method identifies a median RNA secondary structure 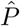 minimizing the sum of distances to a given set **P** = { *P*_1_, …, *P*_*m*_ } of input structures under the Internal-Leafset (IL) or Robinson–Foulds (RF) distance. We used the IL distance, which was reported to perform best in the original publication. From the median structure 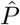, we extracted its set 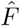 of structural features. Next, to ensure that the `RNAconsensus` solution did not contain any features absent from the input, we took the intersection 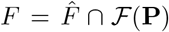 with the set ℱ (**P**) of features present in the input structures **P**. Using these features *F*, we then map the corresponding structural patterns back onto the original input structures to obtain a partition *C*(**P**, *F*) of the structures (following Definition 4). Note that this approach does not allow us to enforce that *C*(**P**, *F*) consists of *τ* clusters.

Second, following Wuchty *et al*. [10], we compute the pairwise symmetric difference of base-pair sets between all RNA secondary structures **P** = {*P*_1_, …, *P*_*m*_}, yielding a distance matrix *D* that quantifies structural dissimilarity.

Using this matrix, the structures are clustered with hierarchical clustering using Ward’s method as the linkage criterion [10]. Next, we cut the resulting dendrogram at the precise height that partitions the data into exactly *τ* clusters. This approach yields a partition ℳof the *m* input structures. Finally, we obtain the corresponding largest set *F* of features corresponding to partition ℳ following Definition 9. We refer to this second baseline approach as `BP-dist + Ward`. Note that unlike `RNAconsensus`, this second approach does respect the *τ* constraint.

## 5 Results

### 5.1 Simulated Data

We used simulations to assess the performance of our three implementations for solving MASS (Problem 1): (i) the integer linear programming formulation (Section 4.1, denoted `MASS-ILP`), (ii) the Max-Subset *τ* -Partitioning exact algorithm (Section 4.2, denoted `MASS-MSTP`), and (iii) the beam search algorithm with beam widths *w* ∈ { 10, 100, 1000 } (Section 4.3, denoted `MASS-BEAM-`*w*). In addition, we included the baseline methods `BP-dist + Ward` [10] and `RNAconsensus` [13] discussed in Section 4.4. We ran these methods on a MacBook Pro M1 Max with 64 GB of RAM and used a time limit of 2 hours per simulation instance.

We generated BMCSP simulation instances in two steps. First, we generated a small binary matrix 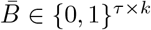 with *τ* unique rows and *k* unique columns. We varied *τ* ∈ { 2, 4, 8, 16 } and varied *k* depending on the choice of *τ* : *k* ∈ { 2, 3 } for *τ* = 2; *k* ∈ { 2, 4 } for *τ* = 4; *k* ∈ { 3, 5 } for *τ* = 8; and *k* ∈ { 4, 6 } for *τ* = 16. Second, we expanded 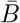 into a *m* × *n* binary matrix *B* with *m* = 20 rows and *n* ∈ { 10, 20, 40, 80 } columns. This entailed duplicating each of the *τ* rows by drawing from relative proportions from a symmetric Dirichlet distribution, followed by randomly generating *n* − *k* additional columns. The entries of each additional column were drawn uniformly at random, but the column was rejected if it matched any of the original *k* columns (or their complement). For each combination of (*τ, k, n, m*), we generated three instances, amounting to a total of 96 simulation instances. As the two baseline methods require input RNA structures, we used Definition 6 to obtain secondary structures **P** from each simulated binary matrix *B*.

We found that the ILP solved all instances to optimality within the 2 hour time limit (Fig. 2a). The MSTP exact algorithm only did so for 87 instances (90.6%). However, for the 87 instances it did solve, its reported solutions matched the ILP solutions in terms of objective value, indicating correctness of both algorithm implementations. While all `MASS-BEAM-`*w* runs were completed within the time limit, the choice of the beam width *w* affected the performance. With decreasing beam widths, the heuristic algorithm ran faster (median of 0.0521 s for *w* = 1000, 0.0283 s for *w* = 100 and 0.0126 s for *w* = 10) at the expense of an increased fraction of non-optimal solutions (1.04% for *w* = 1000, 6.25% for *w* = 100 and 39.6% for *w* = 10). This was also reflected in the ratio | *N* | */* |*N* ^∗^ | of reported optimal columns with a minimum (worst-performance) ratio of 0.750 for *w* = 1000, 0.667 for *w* = 100 and 0.333 for *w* = 10 (Fig. 2c). The two baseline methods `BP-dist + Ward` and `RNAconsensus` were very fast (0.003 s and 1.68 s, respectively) but had worse performance with only 3.13% and 7.29% of instances solved to optimality, respectively. This is unsurprising given that they are not directly optimizing the MASS problem. In particular, `RNAconsensus` has no way of enforcing the presence of at most *τ* clusters (Fig. S5), which allows it to sometimes output a larger number of columns than would be possible when the constraint is enforced (Fig. 2c). The peak memory usage shows a similar trend with `MASS-MSTP` using the most memory, and the heuristic algorithm `MASS-BEAM-`*w* using more memory as *w* increases (Fig. S4). In addition, `ILP` and `RNAconsensus` each used a moderate amount of memory, while `BP-dist + Ward` used the least amount of memory (Fig. S4).

**Figure 2:**
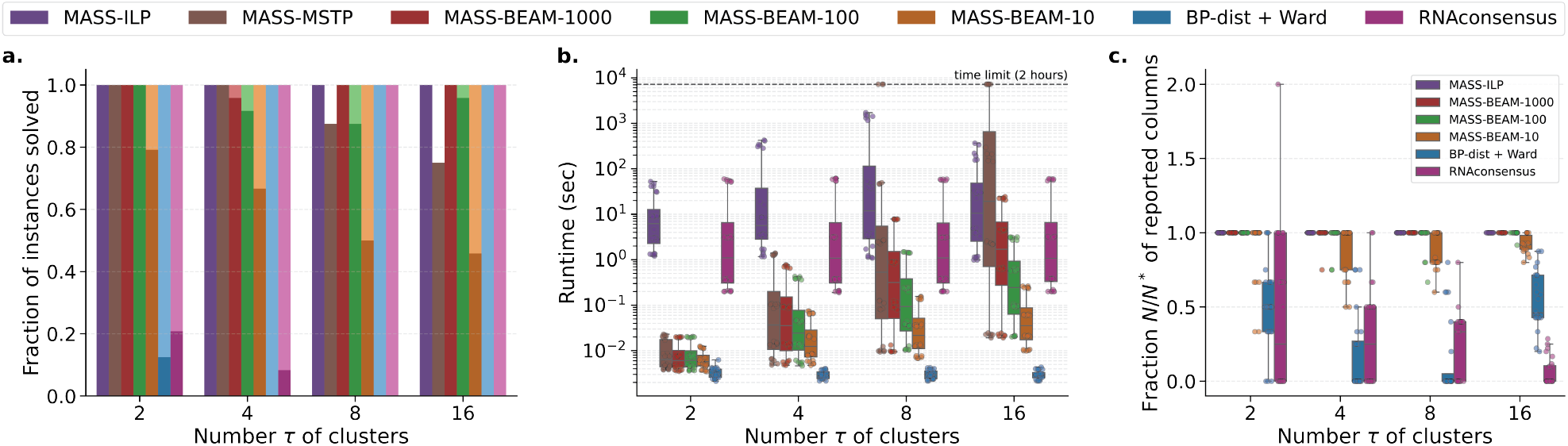
MASS outperforms baseline methods on simulations. (a) Fraction of instances solved within a 2-hour time limit, with shaded stacked bar showing the fraction of instances solved to optimality. (b) Runtime in seconds. (c) Approximation ratio.

In summary, while both `MASS-ILP` and `MASS-MSTP` are exact approaches, the ILP scaled better with increasing *τ* . On the other hand, we found that `MASS-BEAM-`*w* is fast and often returns optimal solutions; this tradeoff between runtime and solution quality can be controlled by the beam width *w*. In contrast, the baseline methods, while fast, consistently failed to find optimal solutions, as they do not optimize the same problem.

### 5.2 CoDNaS-RNA

CoDNaS-RNA [4] is a curated database of experimentally derived structures. Each database entry contains multiple reported structures corresponding to the same RNA sequence. We restricted our analysis to entries whose variants differed only at the secondary-structure level but had identical sequences. As such, we extracted 128 sequence-structure clusters composed of a median of *m* = 4 structures (Fig. 3a), each corresponding to a single RNA sequence ranging from 14 to 3169 nucleotides with a median length of 1500 nucleotides (Fig. 3b). For each instance, we varied the number *τ* of allowed clusters from { 1, …, *m* } . We evaluated how well the `MASS-BEAM-`1000 heuristic and the `BP-dist + Ward` baseline method were able to summarize these RNA structural ensembles. We did not include `RNAconsensus` in this analysis as the number *τ* of clusters could not be specified for this method.

**Figure 3:**
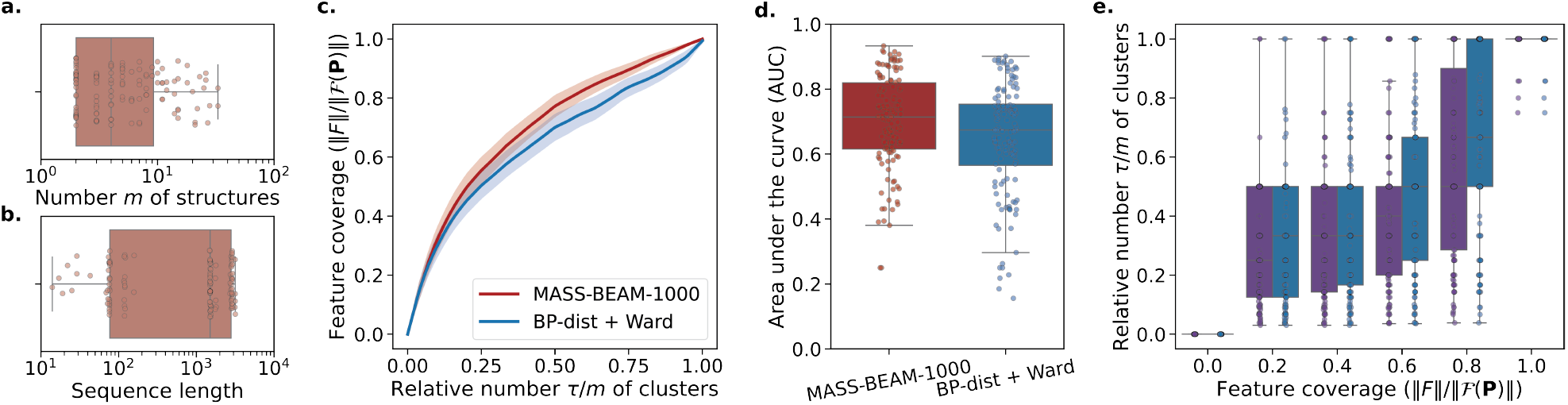
Summarizing 128 CoDNaS–RNA [4] ensembles. (a) Distribution of the number *m* of structures per ensemble. (b) Distribution of sequence lengths in each ensemble. (c) Trade-off between the relative number *τ/m* of clusters and the resulting feature coverage ∥*F* ∥ */* ∥ ℱ (**P**) ∥. Curves show the mean over ensembles for `MASS-BEAM-1000` and the `BP-dist + Ward` baseline, with shaded bands indicating 95% confidence intervals. (d) Area under the *τ/m*–coverage curve (AUC) for each ensemble and method, where larger values indicate more coverage for the range of *τ/m*. (e) Minimum relative number *τ/m* of clusters required to reach a specified coverage cutoff, with lower values indicating that a method attains a given coverage level using fewer clusters.

To compare performance across instances **P** = (*P*_1_, …, *P*_*m*_) with varying number *m* of structures and varying feature space sizes ℱ (**P**), we used the user-specified number *τ/m* of allowed clusters and the *feature coverage* defined as ∥*F* ∥*/* ∥ ℱ (**P**) ∥ where *F* are the identified features by each method. Ideally, a method would simultaneously maximize feature coverage and minimize the relative number of clusters required. To quantify this tradeoff, we computed the area-under-the-curve (AUC), finding that `MASS-BEAM-`1000 achieved a mean AUC value of 0.699 vs. 0.646 for `BP-dist + Ward` (Fig. 3c and boxplots in Fig. 3d). To exemplify, if one wished to achieve a feature coverage of at least 0.800, `MASS-BEAM-`1000 was able to accomplish this with a relative number of 0.500 clusters vs. 0.667 for `BP-dist + Ward` (Fig. 3e). In summary, our method is able to more succinctly (with fewer clusters) and more comprehensively (with high feature coverage) summarize RNA structural ensembles compared to an existing method.

### 5.3 Rfam

For our next experiment, we considered 194 RNA families from Rfam [5], ranging from small families with *m* = 2 structures to large families with as many as *m* = 242 structures (Fig. 4a; median of *m* = 22 structures). Each family consisted of a multiple sequence alignment and a family-level consensus structure that captured the conserved structural motifs characteristic of each family. To obtain secondary structures for the individual sequences within a family, we ran Vienna RNAfold [21] with the consensus structure as a constraint, yielding a collection **P** of aligned structures for each family. In addition, each sequence within a family was annotated by it corresponding species. The selected families had between 2 to 66 species (median: 5; Fig. 4b), with a median of 2 copies of each species. We obtained a ground-truth clustering ℳ^∗^ for each family by grouping sequences according to their species annotations. We compared `MASS-ILP` to the two baseline methods, `BP-dist + Ward` and `RNAconsensus`. While `MASS-ILP` and `BP-dist + Ward` were provided the ground-truth number *τ* = |ℱ ^∗^| of clusters for each family, the `RNAconsensus` method did not allow explicit control of *τ* .

**Figure 4:**
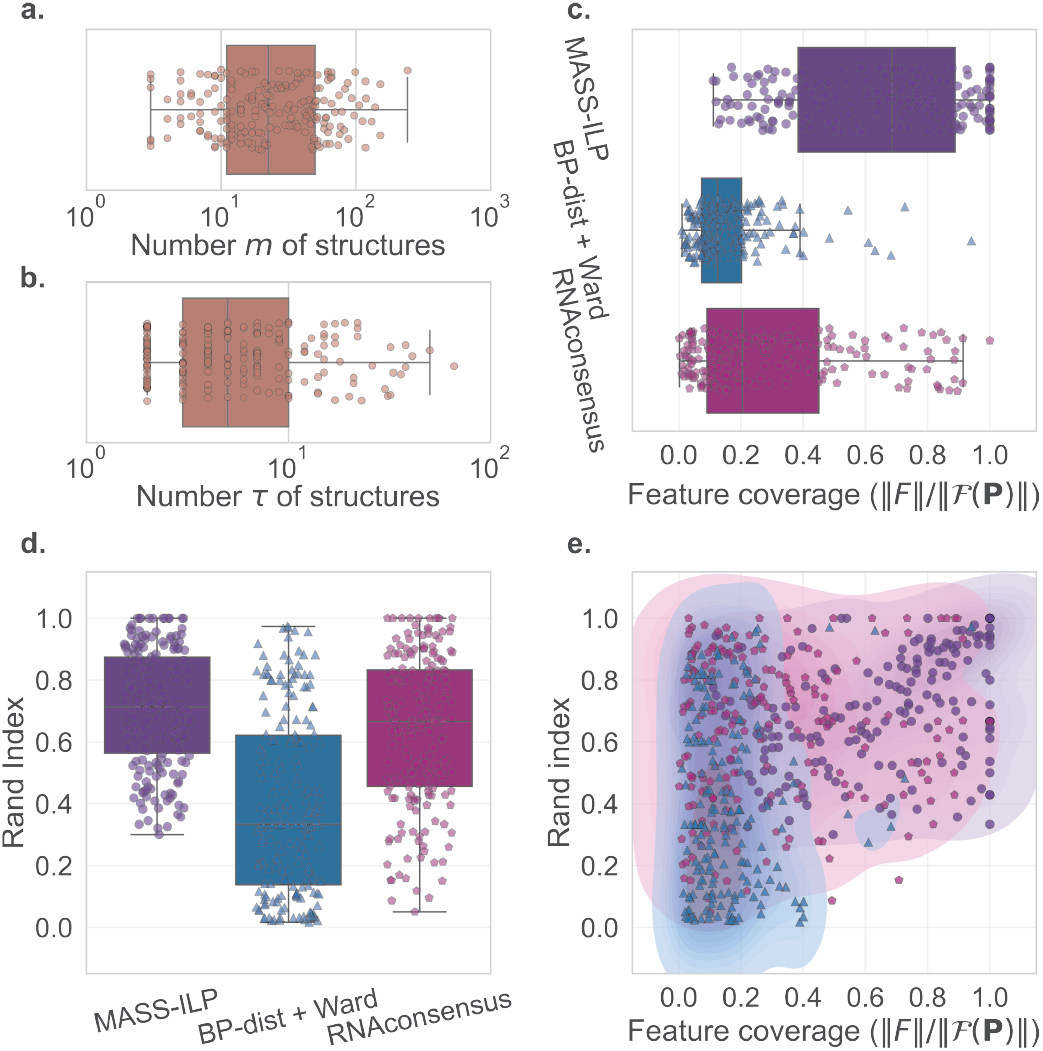
MASS-ILP yielded more accurate summarizations of 194 Rfam families [5]. (a) Distribution of the number *m* of structures per family. (b) Distribution of the number *τ* of species/clusters per family. (c) Feature coverage ∥*F* ∥ */*∥ℱ (**P**) ∥ achieved by each method (d) Rand index of the inferred clusters. (e) Distribution of feature coverage versus Rand index.

Fig. 4c compares the clustering performance of the methods using the Rand index [22], with `MASS-ILP` performing best with a median Rand index of 0.714 for vs. 0.333 for `BP-dist + Ward` and 0.667 for `RNAconsensus`. Fig. 4d reports the corresponding feature coverage ∥*F* ∥*/* ∥ ℱ (**P**) ∥ achieved by the feature set *F* returned by each method. Again `MASS-ILP` performed best (median: 0.686) followed by `RNAconsensus` (median: 0.204) and `BP-dist + Ward` (median: 0.125). Thus, compared to the two baseline methods, `MASS-ILP` yielded more accurate reconstructions of the species-level organization and captured a larger proportion of the shared structural landscape.

### 5.4 mRNA Vaccine Design for SARS-CoV-2 Spike Protein

Finally, we applied `MASS` to summarize *m* = 47 mRNA designs for the SARS-CoV-2 spike protein. These were obtained from the DERNA paper [6], which enumerated sequence-structure pairs (**v**, *P*) encoding for **w** that are Pareto optimal with respect to minimizing the minimum free energy (MFE, a proxy for half-life) and maximizing the codon adaptation index (CAI, a proxy for translation efficiency). Due to the large length of the spike coding sequence (3,819 nucleotides) and therefore the large feature space (10,121 distinct features covering 49,547 loci), running the two exact algorithms was infeasible. Instead, we ran `MASS-BEAM` with a beam width *w* = 1000 for *τ* ∈ { 2, …, 10 } clusters (Fig. 5a).

**Figure 5:**
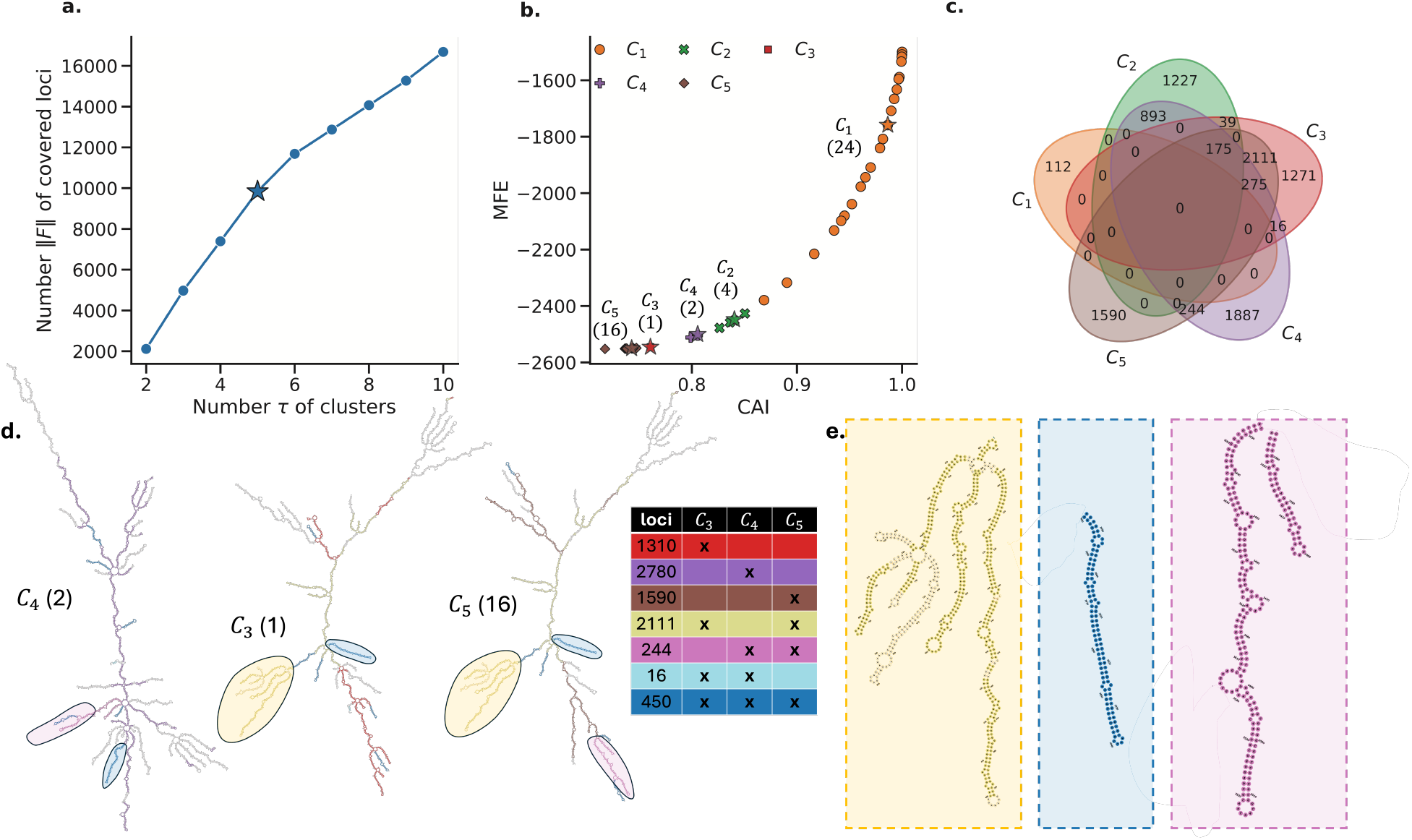
MASS-BEAM-1000 summarizes *m* = 47 alternative mRNA designs encoding for the SARS-CoV-2 spike protein. (a) Number of covered loci as a function of *τ* . (b) Clustering (indicated by shape and color) of the mRNA designs with *τ* = 5 clusters. Star indicates the structure used to represent the cluster in Fig. S6-S10. (c) Number of loci with shared secondary structures among different combinations of clusters. (d) Clusters *C*_3_, *C*_4_, *C*_5_ with coloring indicating shared structural elements among these three clusters (color mapping in (e); number of comprising structures within parentheses). (e) Zoomed in shared structural elements.

We focused our analysis on the solution with *τ* = 5 clusters (Fig. 5b). We found that the largest cluster *C*_1_ (containing 24 structures sharing 112 loci) was the most structurally distinct, sharing no structural elements with the other clusters (Fig. 5c). Further investigation of clusters *C*_3_, *C*_4_, *C*_5_ revealed that *C*_4_ was notably distinct, harboring more unique structural loci (2,780) than either *C*_3_ (1,310) or *C*_5_ (1,590, Fig. 5d). While clusters *C*_3_ and *C*_5_ shared a large set of 2,111 common loci, cluster *C*_4_ shared far fewer features with *C*_3_ (466 loci) and *C*_5_ (694 loci). This suggests that *C*_4_ represents a relatively distinct structural group, whereas *C*_3_ and *C*_5_ are structurally close. In the context of mRNA design, this implies that DERNA undersampled the solution space between the structures in *C*_4_ and those in *C*_3_*/C*_5_. Thus, `MASS` enables the identification of sparsely sampled regions in the design space; targeting these regions for further exploration with DERNA could yield structurally diverse candidates, expanding the pool of potential leads for downstream vaccine validation and development.

## 6 Discussion

In this work, we introduced the Maximum Agreement Secondary Structures (MASS) problem of simultaneously clustering a collection **P** of aligned RNA structures and identifying the structural motifs that characterize those clusters. We demonstrated that existing methods are insufficient, as they only solve half of this combined objective. We proved that MASS is NP-hard and established its equivalence to a constrained binary matrix projection problem. To solve it, we developed three approaches: (i) an exact integer linear program, (ii) an exact combinatorial algorithm, and (iii) an efficient beam-search heuristic adapted from (ii). Our simulations showed that the ILP outperformed the exact combinatorial approach, and that the heuristic algorithm often recovered the optimal solution and provides a tunable speed vs. accuracy tradeoff. Applied to real-world structural ensembles from CoDNaS-RNA, our method significantly outperformed baselines by achieving higher feature coverage with a smaller number of clusters, thus providing a more succinct and comprehensive summary. On Rfam families, our approach yielded more accurate reconstructions of the species-level organization and captured a larger proportion of the shared structural landscape. Finally, our method was able to accurately summarize a solution space of alternative mRNA vaccine designs and identify parts of the solution space that merit further exploration.

There are several future directions. First, our method currently relies on a user-specified number *τ* of clusters. Future work should focus on model selection for *τ* . Second, we could extend the MASS problem to incorporate a balance constraint, requiring a minimum number of structures assigned to each group. Third, we could generalize the problem to allow for soft/approximate matches between the features instead of an exact match. This would increase robustness to minor structural variations within a cluster. Finally, we could incorporate sequence information.

## A Supplementary Proofs

### A.1 Combinatorial Characterization

To prove Proposition 1, we first prove the following helper lemma.

#### Lemma 2.

*Let B* ∈ { 0, 1 } ^*m×n*^ *be a binary matrix obtained from secondary structures* **P** = (*P*_1_, …, *P*_*m*_), *F be a subsets of features* ℱ (*P*) *and N be the corresponding columns of B obtained from F following Definition 5. Then, C*(*F*, **P**) *consists of τ clusters if and only if B*[*N*] *consists of τ rows*.

**(Main Text) Proposition 1**. Let *B* ∈ { 0, 1 } ^*m×n*^ be a binary matrix obtained from secondary structures **P** following Definition 5. Then, there exists an optimal BMCSP solution *N* composed of *k* columns such that *B*[*N*] has *τ* rows if and only if there exists an optimal MASS solution *F* ⊆ ℱ (**P**) covering ∥ *F* ∥ = *k* loci and inducing a partition *C*(*F*, **P**) of *τ* clusters.

*Proof*. (⇒) Let *N* be an optimal BMCSP solution of matrix *B* composed of *k* columns such that *B*[*N*] has *τ* rows. Let *F* be the subset of the feature space ℱ (**P**) corresponding to columns *N*, obtained by reversing the construction. We claim that *F* induces a partition *C*(*F*, **P**) of *τ* clusters and that *F* covers exactly ∥ *F* ∥ = *k* loci. From Lemma 2 it follows that *F* induces a partition *C*(*F*, **P**) of *τ* clusters.

To prove the second part (i.e., ∥ *F* ∥ = *k*), observe that, by Definition 5, we have that each column of *B* corresponds to a locus covered by a feature *f* ∈ ℱ (**P**). Now *N* consists of *k* columns, and *F* is obtained from *N* by reversing the construction in Definition 5. Therefore, *F* must cover at least *k* loci (not necessarily unique). Assume for a contradiction that ∥ *F* ∥ *> k*. In that case, one can obtain a larger subset *N* ^*′*^ ⊋ *N* covering an additional ∥ *F* ∥ − *k >* 0 columns of *B* such that *N* ^*′*^ contains all columns that correspond to features *F* . Recall that we have already shown that *C*(*F*, **P**) consists of *τ* clusters. By Definition 5, we thus have that *B*[*N* ^*′*^] also consists of *τ* rows. This would imply that *N* consisting of |*N* | = *k <* |*N* ^*′*^| columns is not optimal — a contradiction. Hence, *F* covers exactly ∥*F* ∥ = *k* loci.

It remains to show that *F* is an optimal MASS solution. Assume for a contradiction that there exists another feasible solution *F* ^*′*^ covering more than ∥ *F* ^*′*^ ∥ *>* ∥ *F* ∥ = | *N* | = *k* loci and inducing a partition *C*(*F* ^*′*^, **P**) of *τ* ^*′*^≤ *τ* clusters. Let *N* ^*′*^ the columns corresponding to features *F* ^*′*^. We thus have | *N* ^*′*^ | = ∥ *F* ^*′*^ ∥ *>* ∥ *F* ∥ = | *N* | = *k*. From Lemma 2 it follows that the number of rows *B*[*N* ^*′*^] equals the number | *C*(**P**, *F* ^*′*^) | = *τ* ^*′*^ of clusters. Since | *C*(**P**, *F* ^*′*^) | = *τ* ^*′*^ ≤ *τ* and | *N* ^*′*^ | *>* | *N* |, we have that *N* is not an optimal BMCSP solution, a contradiction. Hence, *F* is an optimal MASS solution.

(⇐) Let *F* ⊆ ℱ (**P**) be an optimal MASS solution covering ∥*F* ∥ = *k* loci and inducing a partition *C*(*F*, **P**) of *τ* clusters. Moreover, let *B* be the binary matrix obtained from **P**. We must show that there exists an optimal BMCSP solution *N* for matrix *B* such that *B*[*N*] has *τ* rows and *B*[*N*] has *k* columns (i.e., | *N* | = *k*). Let *N* be obtained from *F* by reversing the construction of *B* from **P** following Definition 5. From Lemma 2 it follows that the number of rows *B*[*N*] equals the number | *C*(**P**, *F*) | = *τ* of clusters. By construction of *N* following Definition 5, we have | *N* | = ∥ *F* ∥ = *k*.

It remains to show that *N* is an optimal BMCSP solution. Assume for a contradiction there exists another solution *N* ^*′*^ such that *B*[*N* ^*′*^] contains *τ* ^*′*^ ≤ *τ* rows and | *N* ^*′*^ | *>* | *N* | = *k*. Let *F* ^*′*^ ⊆ ℱ (**P**) be the features obtained from *N* ^*′*^. By Definition 5, we have that ∥ *F* ∥ ≥ ∥ *N* ^*′*^ ∥ . Thus, we have that ∥ *F* ^*′*^ ∥ ≥ ∥*N* ^*′*^ ∥ *>* | *N* | = ∥ *F* ∥ = *k*. Moreover, by Lemma 2, it follows that *C*(*F* ^*′*^, **P**) consists of *τ* ^*′*^ ≤ *τ* clusters given that *B*[*N* ^*′*^] consists of *τ* ^*′*^ ≤ *τ* rows. Thus, *F* ^*′*^ is a feasible solution to MASS instance (**P**, *τ*) and therefore *F* is not an optimal MASS solution; a contradiction. Hence, *F* must be an optimal solution to the MASS instance (**P**, *τ*).

To prove Proposition 2, we prove three helper lemmas. First, we show that we may assume without loss of generality that *B* does not contain any all 0s columns.

#### Lemma 3.

*Let B*^*′*^ ∈ {0, 1}^*m×n*^*′ be the matrix obtained from B* ∈ {0, 1}^*m×n*^ *after removal of columns X* ⊆ [*n*] *composed of only* 0*-entries, let N* ^*′*^ ⊆ [*n*^*′*^] *and τ* ^*′*^ ∈ [*m*]. *Then N* ^*′*^ *induces a binary matrix B*[*N* ^*′*^] *of τ* ^*′*^ *rows if and only if N* ^*′*^ ∪ *X induces a binary matrix B*[*N* ^*′*^ ∪ *X*] *of τ* ^*′*^ *rows*.

*Proof*. This follows from the premise, implying that all *m* rows of *B* are identical in columns *X*. Next, the use of the BP representation implies the following lemma.

#### Lemma 4.

*Let* **P** *be the* MASS-BP *instance obtained from binary matrix B following Definition 6 and F* ⊆ *F*_bp_(**P**) *be a feasible* MASS-BP *solution. Then, the number* ∥*F* ∥ *of covered loci is even*.

*Proof*. As the feature space ℱ_bp_(**P**) resulting from this reduction uses the BP representation (Definition 1), each subset of ℱ_bp_(**P**) will cover an even number of loci.

Finally, we have the following lemma.

#### Lemma 5.

*Let* **P** *be the secondary structures obtained from a binary matrix B, N be a subset of columns of B and F the corresponding subset of features* ℱ_bp_(**P**) *following Definition 6. Then, B*[*N*] *consists of τ rows if and only if C*(*F*, **P**) *consists of τ clusters*.

*Proof*. By Definition 4, two structures *P*_*i*_, *P*_*j*_ fall into the same cluster of *C*(**P**, *F*) if and only if *f* (*P*_*i*_) ∩ *F* = *f* (*P*_*j*_) ∩ *F* . By Definition 6, two rows *i* and *j* of *B*[*N*] are identical if and only if *f* (*P*_*i*_) ∩ *F* = *f* (*P*_*j*_) ∩ *F* . Therefore, two structures *P*_*i*_, *P*_*j*_ are in the same cluster of *C*(**P**, *F*) if and only if their corresponding rows *i* and *j* of *B*[*N*] are identical. Hence, the number of rows of *B*[*N*] equals |*C*(**P**, *F*)| = *τ* .

**(Main Text) Proposition 2**. Let **P** be the MASS-BP instance obtained from binary matrix *B* (without any all 0s columns) following Definition 6. Then, there exists an optimal MASS-BP solution *F* ⊆ ℱ_bp_(**P**) covering ∥ *F* ∥ = 2*k* loci and inducing |*C*(*F*, **P**) | = *τ* clusters if and only if there exists an optimal BMCSP solution *N* such that *B*[*N*] has *k* columns and *τ* rows.

*Proof*. (⇒) Let *F* be an optimal MASS-BP solution of secondary structures **P** covering ∥ *F* ∥ = 2*k* loci and inducing | *C*(*F*, **P**) | = *τ* clusters. Let *N* be a subset of columns of matrix *B* corresponding to *F* by reversing the construction described in Definition 6. We must show that *B*[*N*] has *k* columns and *τ* rows. In the BP representation (Definition 1), each feature *f* ∈ ℱ_bp_(**P**) covers exactly two loci. As ∥ *F* ∥ = 2*k*, we have that *F* consists of *k* features. By construction following Definition 6, we thus have that *N* = | *F* | = *k* and thus *B*[*N*] consists of *k* columns. It follows from Lemma 5 that *B*[*N*] consists of *τ* rows.

It remains to show that *N* is an optimal BMCSP solution, which we prove by contradiction. Let *N* ^*′*^ be a feasible solution to BMCSP instance (*B, τ*) such that *B*[*N* ^*′*^] consists of | *N* ^*′*^ | *>* | *N* | = *k* columns and *τ* ^*′*^ ≤ *τ* rows. We reverse the construction of matrix *B* (following Definition 6) to obtain the feature subset *F* ^*′*^ corresponding to *N* ^*′*^. By construction following Definition 6 and Lemma 4, we have | *N* ^*′*^ | = ∥ *F* ^*′*^ ∥ */*2 *> k/*2. By Lemma 5, *C*(*F* ^*′*^, **P**) consists of *τ* ^*′*^ clusters. This means that *F* ^*′*^ is a feasible solution to BMCSP instance **P** inducing | *C*(*F*, **P**) | = *τ* ^*′*^ ≤ *τ* clusters and | *N* ^*′*^ | = ∥ *F* ^*′*^ ∥ */*2 *> k/*2 — implying *F* was not an optimal solution, a contradiction.

(⇐) Let *N* be an optimal BMCSP solution inducing a matrix *B*[*N*] composed *τ* rows and 2*k* columns. Let *F* be the corresponding subset of features ℱ_bp_(**P**) obtained by reversing the construction of Definition 6. We must show that *C*(*F*, **P**) consists of *τ* clusters and the number ∥ *F* ∥ of covered loci equals *k*. By Lemma 5, *C*(*F*, **P**) consists of *τ* clusters. By the premise, we know that *B* does not contain any all 0s columns. Therefore each column in *N* corresponds to exactly one base pairing covering two loci. Since *N* consists of *k* columns, we have ∥ *F* ∥ = 2*k*.

It remains to show that *F* is an optimal MASS-BP solution. Assume for a contradiction there exists another feasible MASS-BP solution *F* ^*′*^ such that | *C*(*F* ^*′*^, **P**) | = *τ* ^*′*^ ≤ *τ* and ∥ *F* ^*′*^ ∥ *>* ∥*F* ∥ = 2*k*. Let *N* ^*′*^ be the subset of columns of *B* corresponding to *F* ^*′*^ reversing the construction of Definition 6. By Lemma 5, we have *B*[*N* ^*′*^] consists of *τ* ^*′*^ rows. Moreover, as each column of *B* corresponds to a single base pairing, *F* ^*′*^ covers |*N* ^*′*^|*/*2 loci, i.e., ∥*F* ^*′*^∥ = |*N* ^*′*^|*/*2 *> k/*2. This means we have a contradiction regarding the optimal of *N* .

**(Main Text) Theorem 1**. BMCSP is equivalent to MASS.

*Proof*. This directly follows from Proposition 1 and Proposition 2.

### A.2 Complexity

**(Main Text) Lemma 1**. Let (*B, τ*) be an instance of BMCSP obtained from a Independent Set instance (*G, k*) following Definition 7. Then, there exists an independent set *I* of *G* composed of |*I*| = *k* vertices if and only if there exists a subset *N* of columns inducing a submatrix *B*[*N*] composed of |*N* | = *k* columns and at most *τ* = *k* + 1 rows.

*Proof*. (⇒) Let *I* be an independent set of *G* of size |*I* | = *k*. Let *N* be the corresponding set of column indices of matrix *B* following the construction of Definition 7. Since no two vertices in *I* are adjacent, for each edge *e*_*i*_ = (*v*_*j*_, *v*_*j*_*′*) at most one of { *j, j*^*′*^ } belongs to *N* . If neither is chosen, the three corresponding rows { 3(*i* − 1)+1, 3(*i* − 1)+ 2, 3(*i* − 1) + 3 } of *B*[*N*] collapse to the all-zero row vector. If exactly one is chosen, say *j* w.l.o.g., they collapse to two row vectors: one vector of all-zeros and one vector composed of exactly one 1 at column *j* and 0 entries elsewhere. Because no edge has both endpoints in *I*, the projection never yields all three distinct row patterns. Consequently, every row of *B*[*N*] is either the all-zero vector or a vector with a single 1 in one of the chosen columns. The number *τ* of distinct rows of *B*[*N*] is therefore at most *k* + 1, meeting the constraint.

(⇐) Let *N* be a subset of columns of *B* inducing a submatrix *B*[*N*] composed of |*N*| = *k* columns and at most *τ* = *k* + 1 rows. Let *I* be the subset of vertices corresponding to *N* following Definition 7. We start by observing that each column *j* of *N* contributes to rows to *B*[*N*], a row composed of a 1 at column *j* and 0 entries elsewhere and an all-zero row. This mean that the *N* = *k* vertices induce at least *k* + 1 distinct rows in *B*[*N*]. Now, if *N* were to contain two adjacent vertices (*v*_*j*_, *v*_*j*_*′*) then the three rows corresponding to edge *e*_*i*_ = (*v*_*j*_, *v*_*j*_*′*) would project to (1, 0), (0, 1), and (1, 1) — the latter being an additional row in *B*[*N*] not accounted for previously. This would mean that *B*[*N*] must contain strictly more than *k* + 1 distinct row, a contradiction. Hence no two vertices in *I* are adjacent, and *I* forms an independent set of size *k*.

**(Main Text) Theorem 2**. MASS is NP-hard.

*Proof*. This follows Theorem 1, Theorem 3 and the fact that the two reductions take polynomial time.

**(Main Text) Theorem 3**. BMCSP is NP-hard.

*Proof*. This follows from Lemma 1 and the fact that the reduction takes polynomial time.

### A.3 Correctness of Algorithms

**(Main Text) Proposition 3**. Let *N* ^∗^ = *N* (ℳ) be the ℳ-induced column set of a given partition ℳ of rows [*m*] of matrix *B* ∈ *{*0, 1*}*^*m×n*^. Then, for every subset *N* ^*′*^ ⊆ [*n*] of columns such that ℳ (*N* ^*′*^) equals ℳ it holds that *N* ^*′*^ ⊆ *N* ^∗^.

*Proof*. We prove this by contradiction. Let *N* ^*′*^ ⊆ [*n*] be a subset of columns such that *N* ^*′*^ *\ N* ≠ ∅. Let column *j*be in *N* ^*′*^ but not in *N* . Since ℳ (*N* ^*′*^) = ℳ, it holds that, for each row set *M*_*i*_ ∈ ℳ, column *j* must consist of either 0 or 1 entries. However, by Definition 9, *j* must be in *N* ^∗^ = *N* (ℳ), a contradiction.

**(Main Text) Theorem 4**. Let ℳ = *{M*_1_, …, *M*_*τ*_ *}* be a partition of rows [*m*] of matrix *B* ∈ *{*0, 1*}*^*m×n*^ such that there exists a subset *N* ^*′′*^ ⊆ [*n*] of columns where ℳ (*N* ^*′′*^) = ℳ. Then, there exists a subset *N* ^*′*^ ⊆ *N* ^*′′*^ ⊆ [*n*] of columns of size |*N* ^*′*^| ≤ *τ* − 1 such that *N* ^*′*^ also induces partition ℳ, i.e. ℳ (*N* ^*′*^) = ℳ.

*Proof*. We construct *N* ^*′*^ iteratively, maintaining *N* ^*′*^ *⊆N* ^*′′*^ as an invariant. First, we initialize ℓ = 1 and *N* ^*′*^ =∅ such that the induced partition ℳ (*N* ^*′*^) has ℓ = 1 part, which is [*m*]. Note that we respect the invariant by definition, i.e. *N* ^*′*^ = ∅ ⊆ *N* ^*′′*^. Next, while ℓ *< τ*, we extend *N* ^*′*^ with a single column as follows: Since *N* ^*′*^ is a subset of *N* ^*′′*^ and *N* ^*′*^ has ℓ *< τ* parts, partition ℳ (*N* ^*′*^) must be coarser than ℳ. This means there exists at least one part *M* ^*′*^ *∈ ℳ* (*N* ^*′*^) that contains two or more parts from ℳ. Let *M*_1_ and *M*_2_ be two distinct parts from ℳ such that *M*_1_ ∪ *M*_2_ ⊆ *M* ^*′*^. This implies that rows *M*_1_ and *M*_2_ of *B* are identical when restricted to the columns in *N* ^*′*^. However, we know that the same two sets *M*_1_, *M*_2_ of rows of *B* differ over *N* ^*′′*^. Therefore, there must exist a column *j* ∈ *N* ^*′′*^*\N* ^*′*^ that distinguishes the two row sets *M*_1_ and *M*_2_. We add this column *j* to our set: *N* ^*′*^ *→N* ^*′*^∪*{j}* . This new column *j* splits the part *M* ^*′*^ into at least two new parts (one for rows with 0 at *j*, one for rows with 1 at *j*). This guarantees that *M*_1_ and *M*_2_ are now in separate parts. The new partition ℳ (*N* ^*′*^) is a strict refinement of the previous partition, and the number ℓ = |ℳ (*N* ^*′*^)| of parts increases by at least 1, but ℓ *≤ τ* as ℳ (*N* ^*′′*^) =ℳ . The loop starts with ℓ = 1. Each iteration adds exactly one column and increases ℓ by at least 1. To get from ℓ = 1 to ℓ = *τ*, we need at most *τ* − 1 iterations. Therefore, the final set *N* ^*′*^ will have a size |*N* ^*′*^| ≤ *τ* − 1. By the end of the process, we have ℳ (*N* ^*′*^) = ℳ.

## B Supplementary Methods

We provide pseudocode for the following algorithms discussed in the main text.

- Algorithm 1: The *O*(*mnτ*^*m*^) ExhaustivePartitionSearch exact algorithm.
- Algorithm 2: The *O*(*mn*^*τ*^) ExhaustiveColumnSubsetSearch exact algorithm.
- Algorithm 3: The *O*(*mn*^*τ*^) MSTP exact algorithm.
- Algorithm 4: The *O*(*τwmn*^2^) MSTP-Beam heuristic algorithm.

### Algorithm 1

ExhaustivePartitionSearch(*B, τ*)

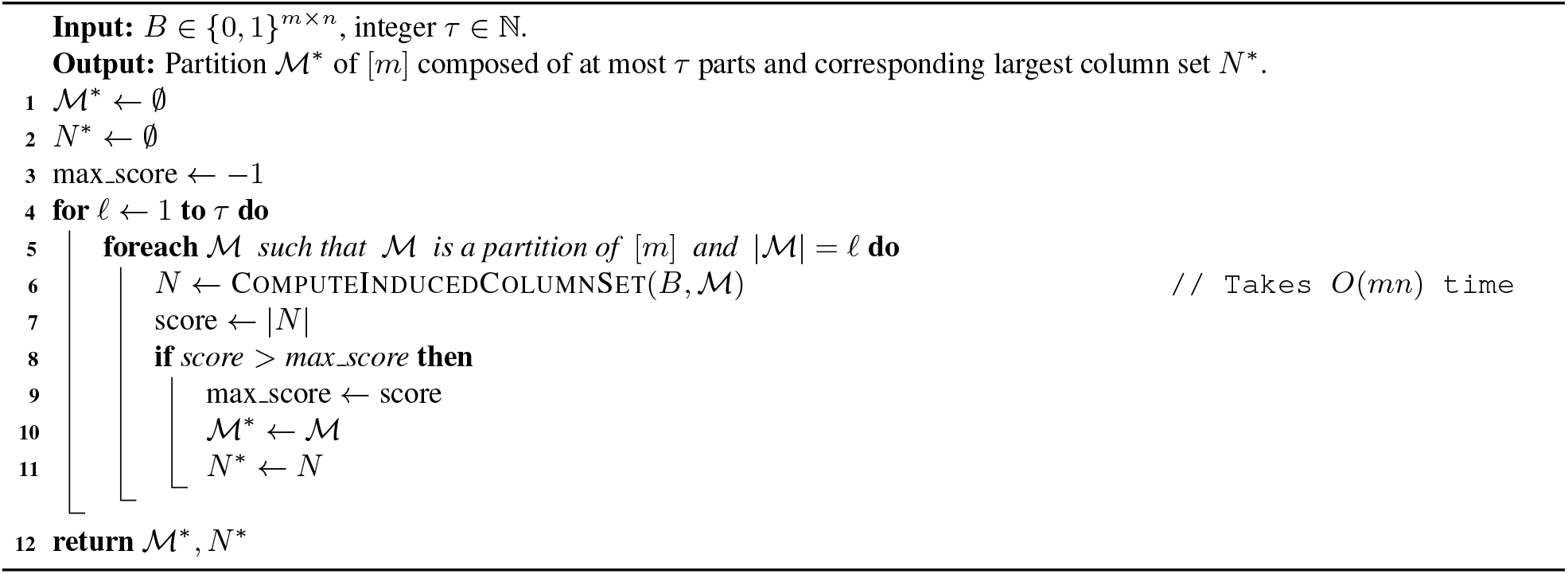

### Algorithm 2

ExhaustiveColumnSubsetSearch(*B, τ*)

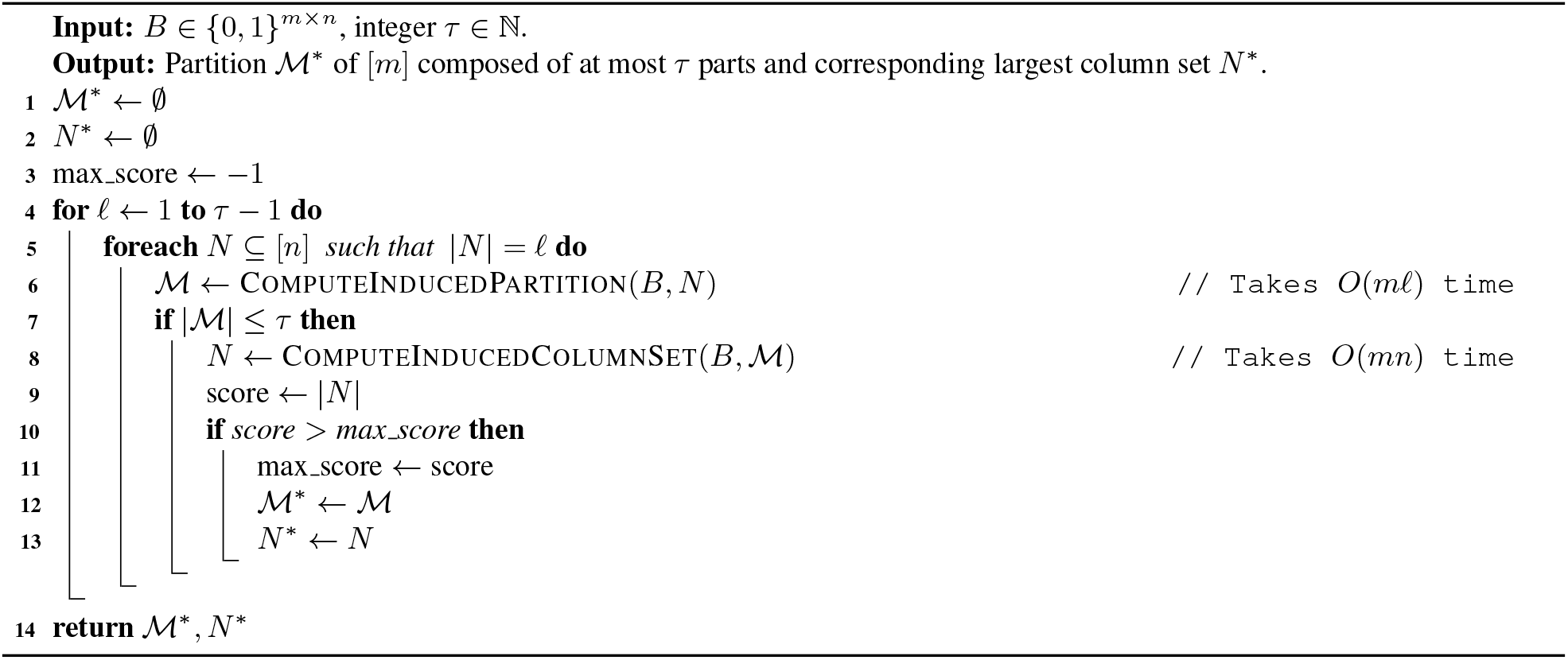

## C Supplementary Figures

### Algorithm 3

MSTP(*B, τ*)

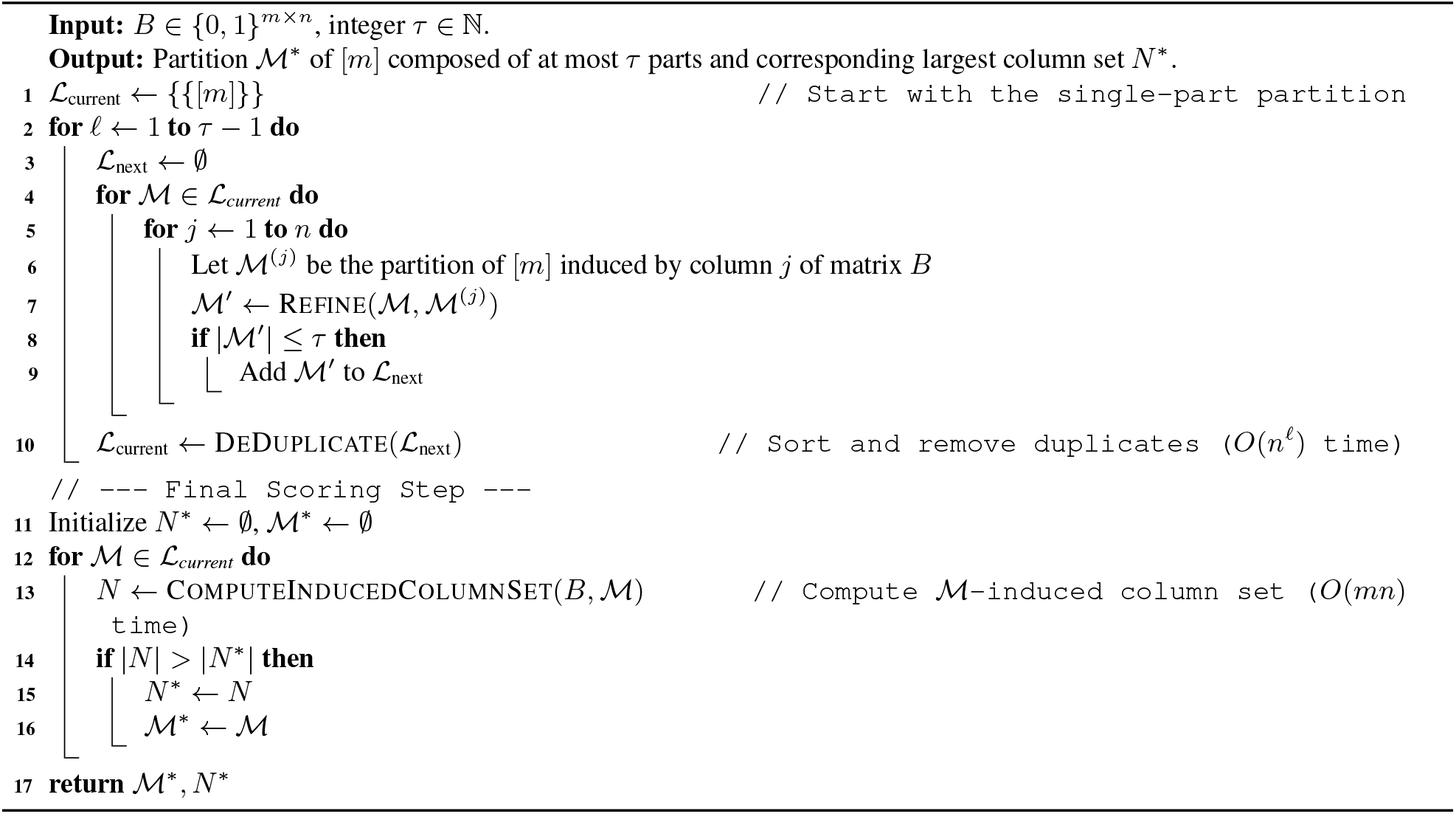

### Algorithm 4

MSTP-Beam(*B, τ, w*)

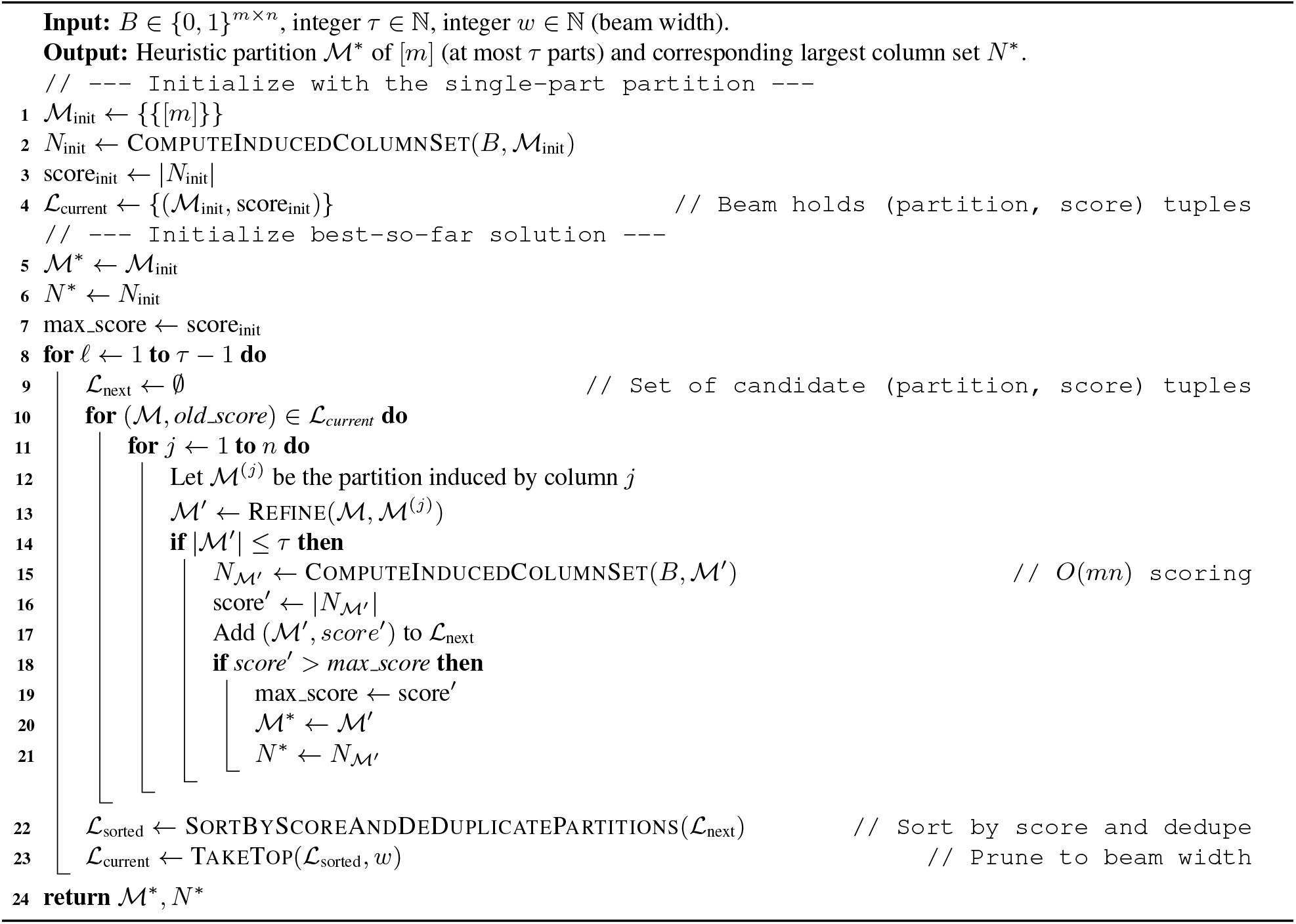

**Figure S1:**
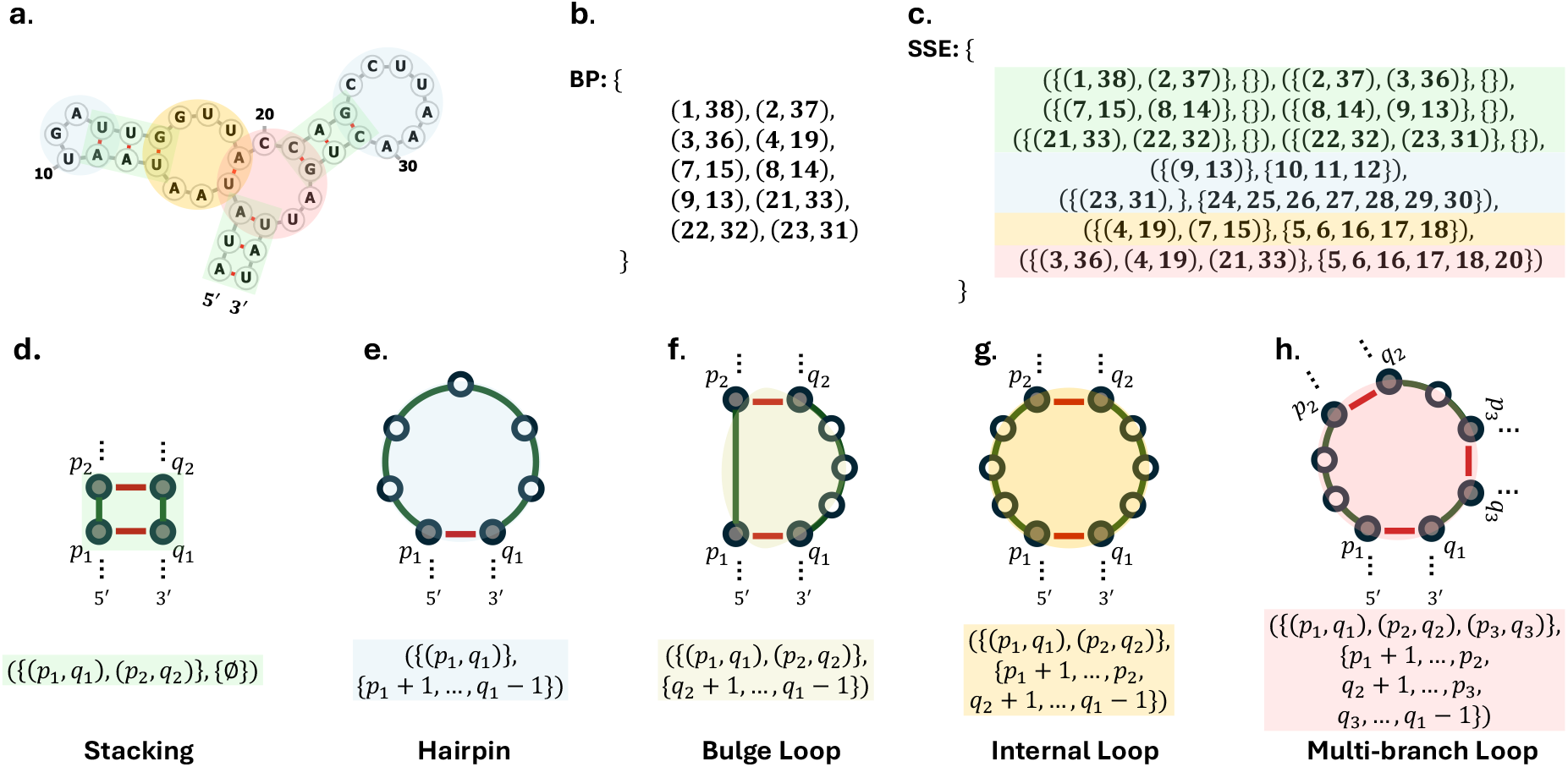
Secondary structure representations. (a) Example secondary structure, with coloring indicating structural elements depicted in (d-h). (b) The base pairing (BP) representation (Definition 1). (c) The secondary structure element (SSE) representation (Definition 2). (d-h) Secondary structural elements that arise in pseudoknot-free structures.

**Figure S2:**
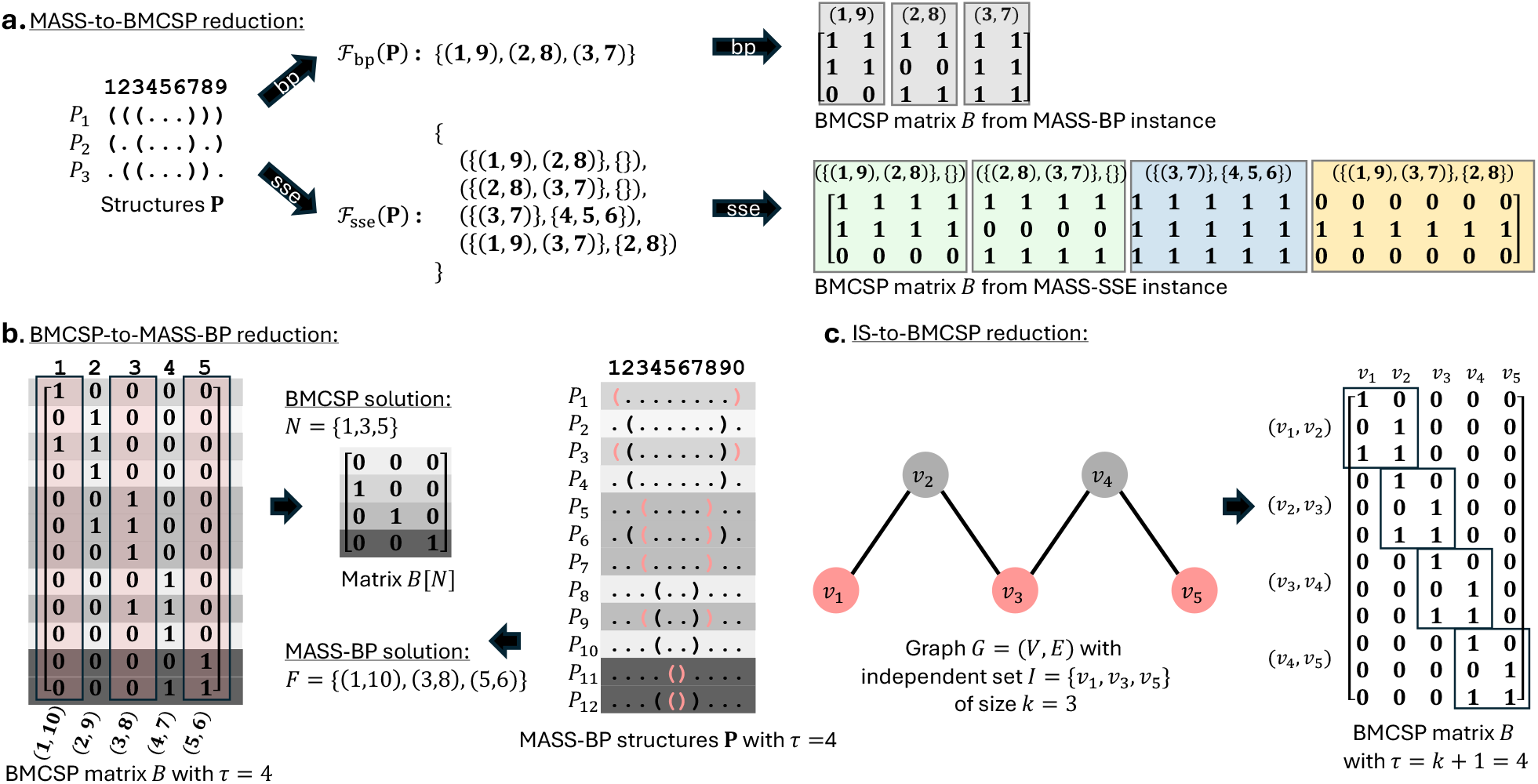
MASS and BMCSP are equivalent, and Independent Set reduces to the latter. (a) Reduction, as in Definition 5, from MASS (Problem 1) to BMCSP (Problem 2) using either the base pairing (BP, Definition 1) or the secondary structure element (SSE, Definition 2 and color coding given in Fig. S1) representation. (b) Reduction, as in Definition 6, from BMCSP to MASS-BP with *τ* = 4. We also illustrate the optimal MASS-BP and corresponding BMCSP solution. (c) Reduction, as in Definition 7, from Independent Set (IS, Problem 3) to BMCSP. The given graph *G* contains an independent set *I* = *{v*_1_, *v*_3_, *v*_5_*}* of size *k* = 3, which corresponds to the optimal BMCSP solution *N* = *{*1, 3, 5*}* with *τ* = *k* + 1 = 4 induced rows in *B*[*N*] shown in (b).

**Figure S3:**
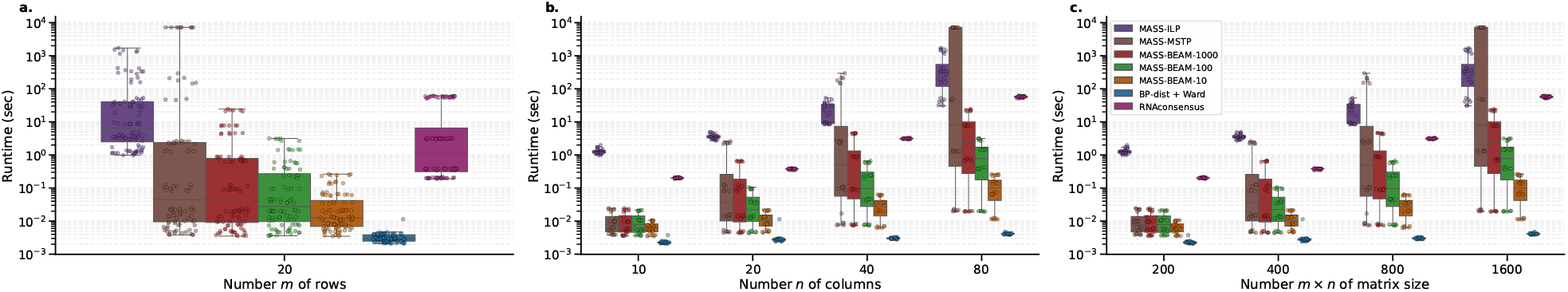
Runtime of simulations. (a) By number *m* of rows. (b) By number *n* of columns. (c) By number *m × n* of entries.

**Figure S4:**
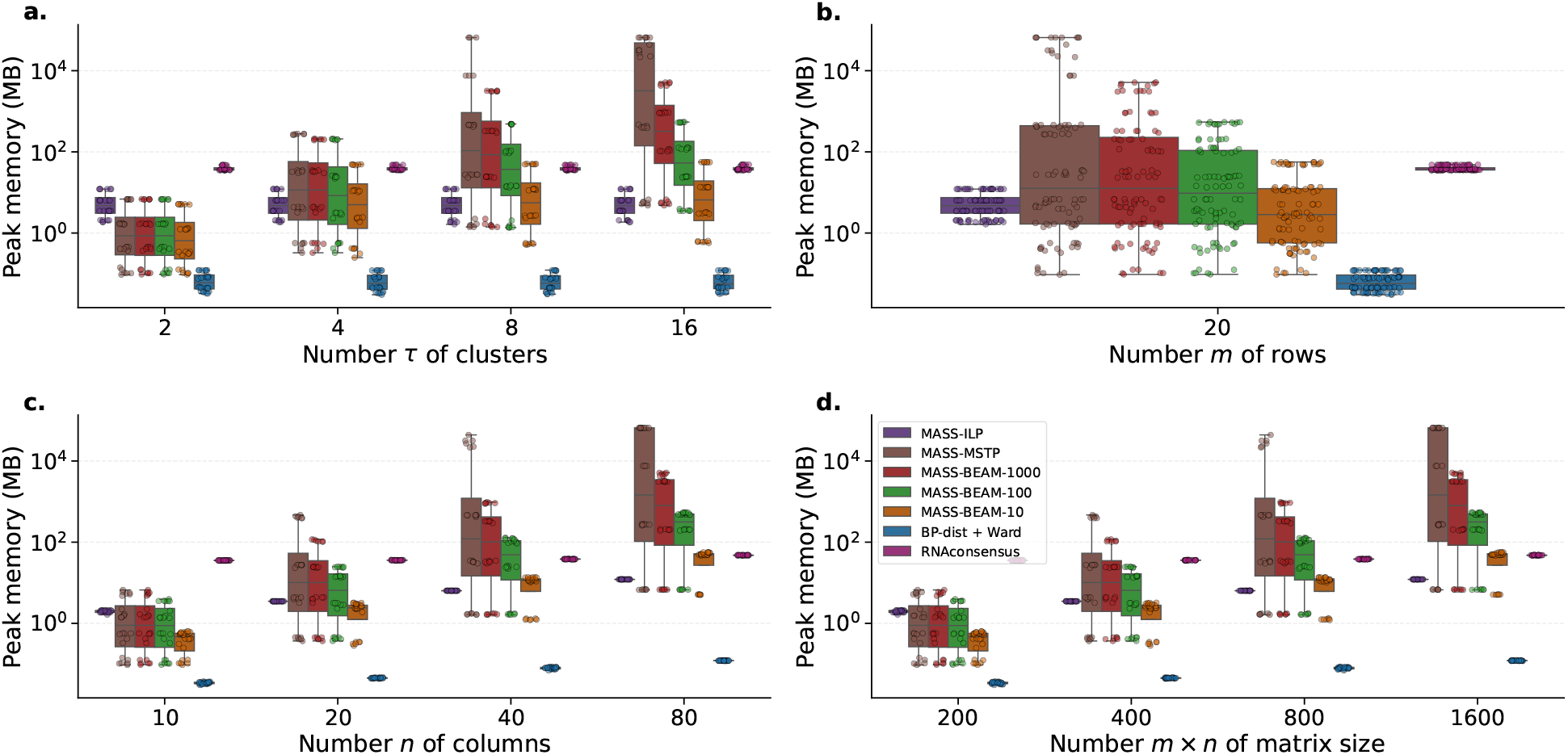
Peak memory usage of simulations. (a) By number *τ* of clusters. (b) By number *m* of rows. (c) By number *n* of columns. (d) By number *m × n* of entries.

**Figure S5:**
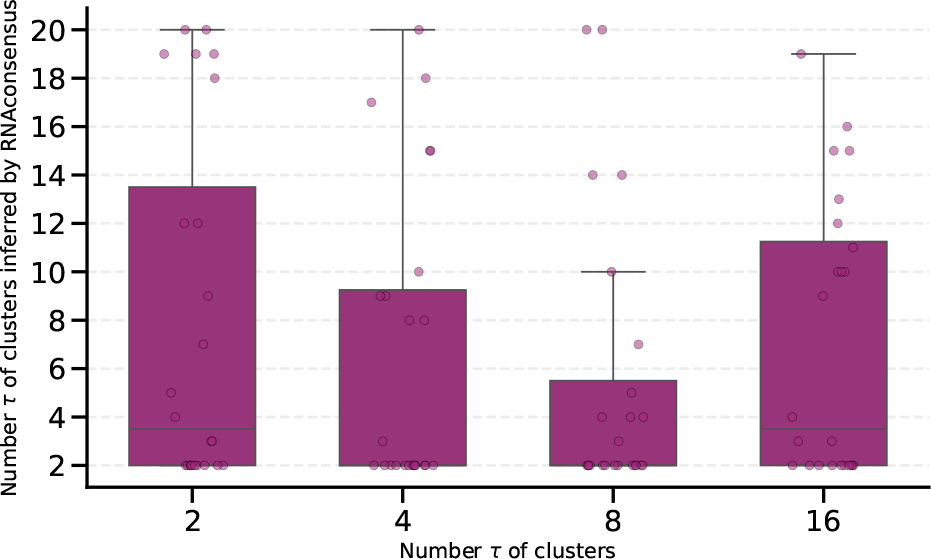
RNAconsensus does not respect the *τ* constraint. *x*-axis shows the ground-truth number of clusters whereas *y*-axis shows the number of clusters inferred by `RNAconsensus`.

**Figure S6:**
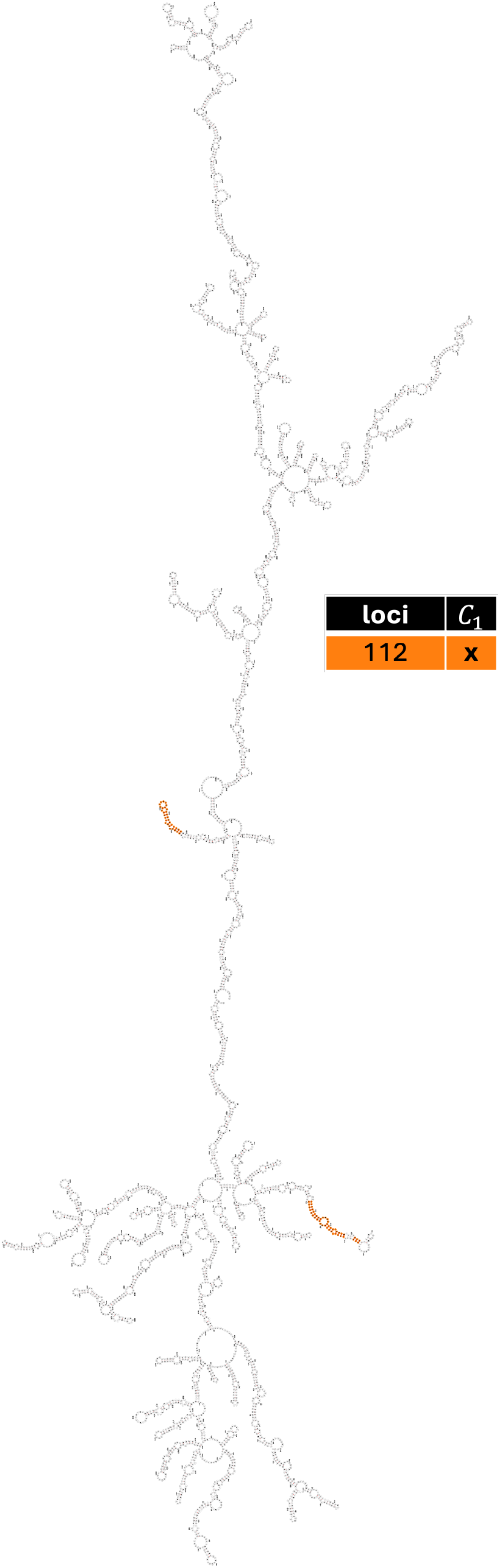
Cluster *C*_1_ of spike mRNA designs. Cluster *C*_1_ contains 24 structures, with CAI ranging from 0.869 to 1.0 and MFE from −2379.2 kcal/mol to −1499.3 kcal/mol. Table and coloring indicates cluster membership of loci with shared secondary structure.

**Figure S7:**
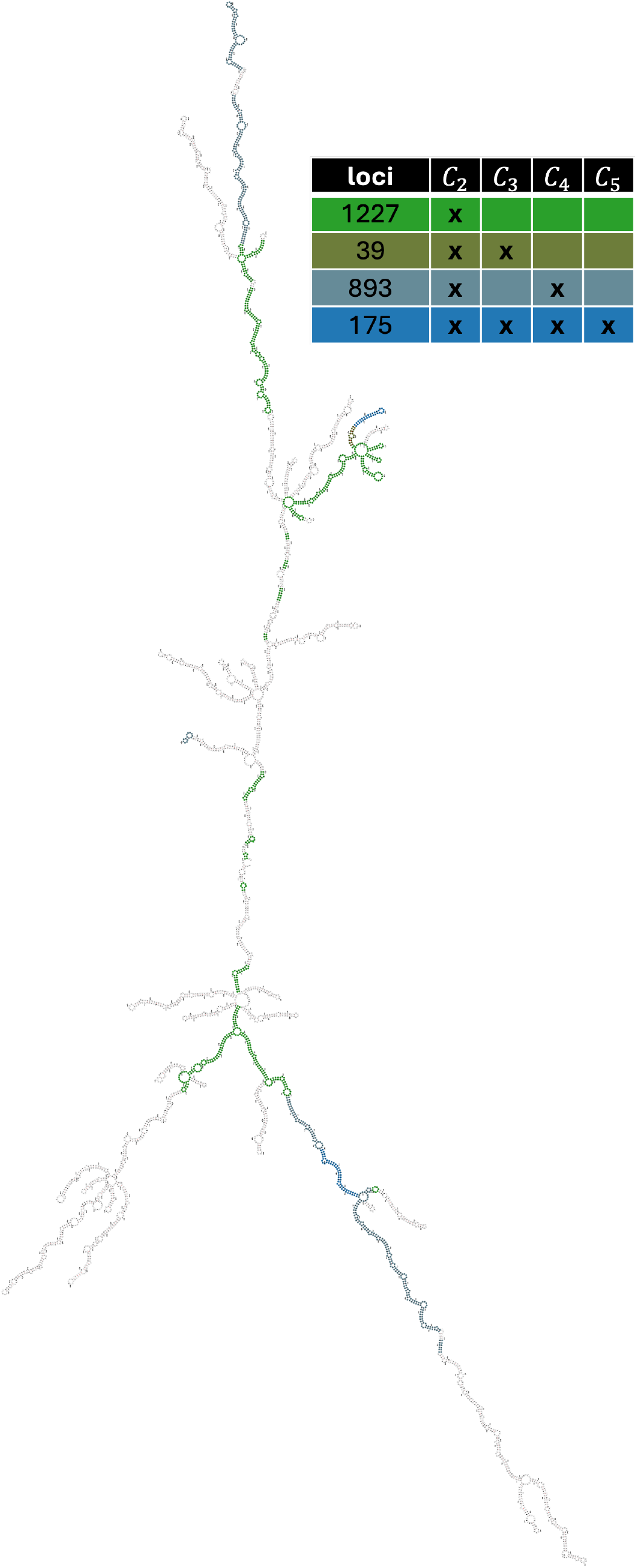
Cluster *C*_2_ of spike mRNA designs. Cluster *C*_2_ contains 4 structures, with CAI ranging from 0.826 to 0.850 and MFE from −2477.6 kcal/mol to −2426.5 kcal/mol. Table and coloring indicates cluster membership of loci with shared secondary structure

**Figure S8:**
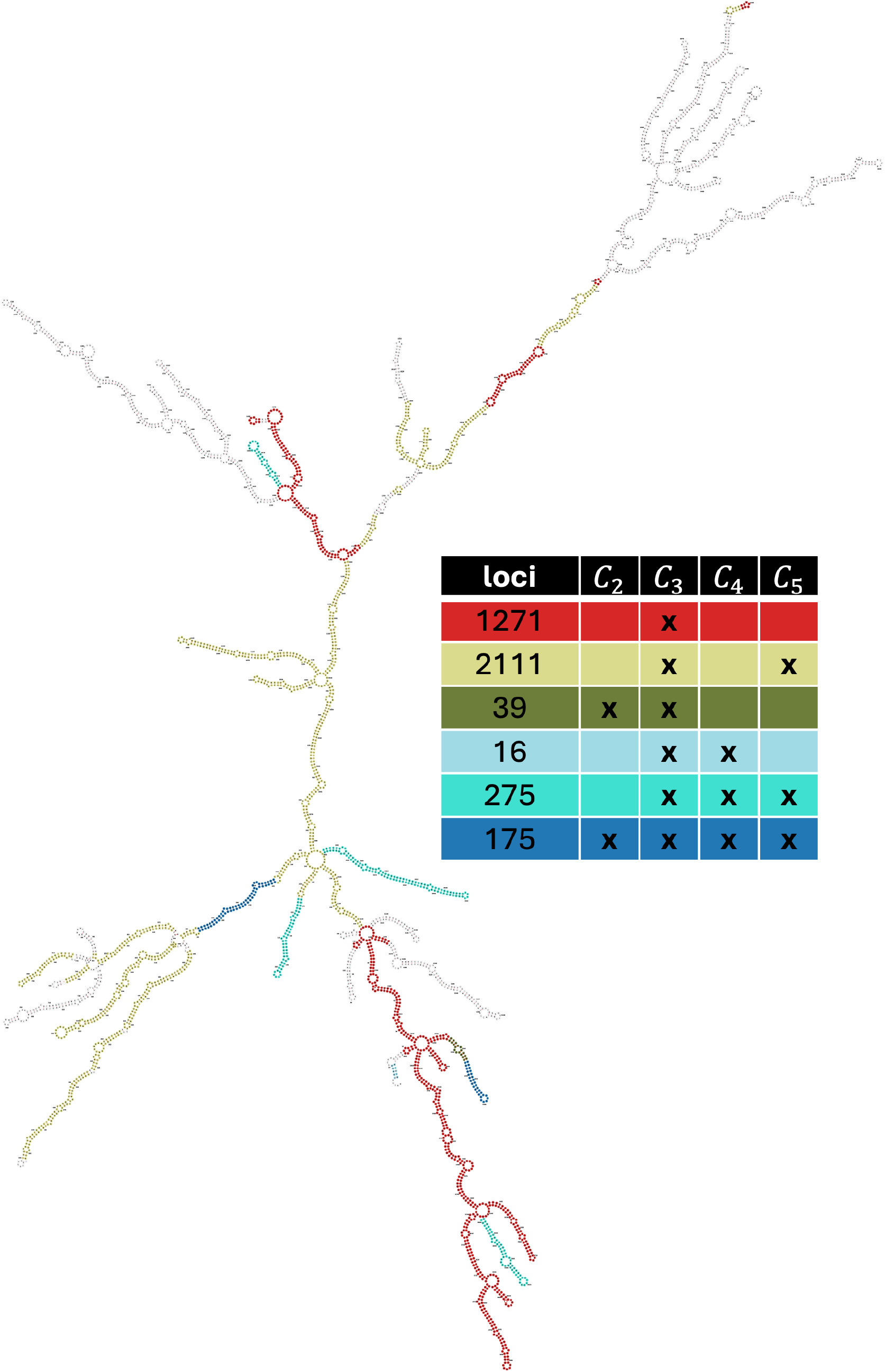
Cluster *C*_3_ of spike mRNA designs. Cluster *C*_3_ contains 1 structures, with CAI of 0.760 and MFE of −2545.5 kcal/mol. Table and coloring indicates cluster membership of loci with shared secondary structure.

**Figure S9:**
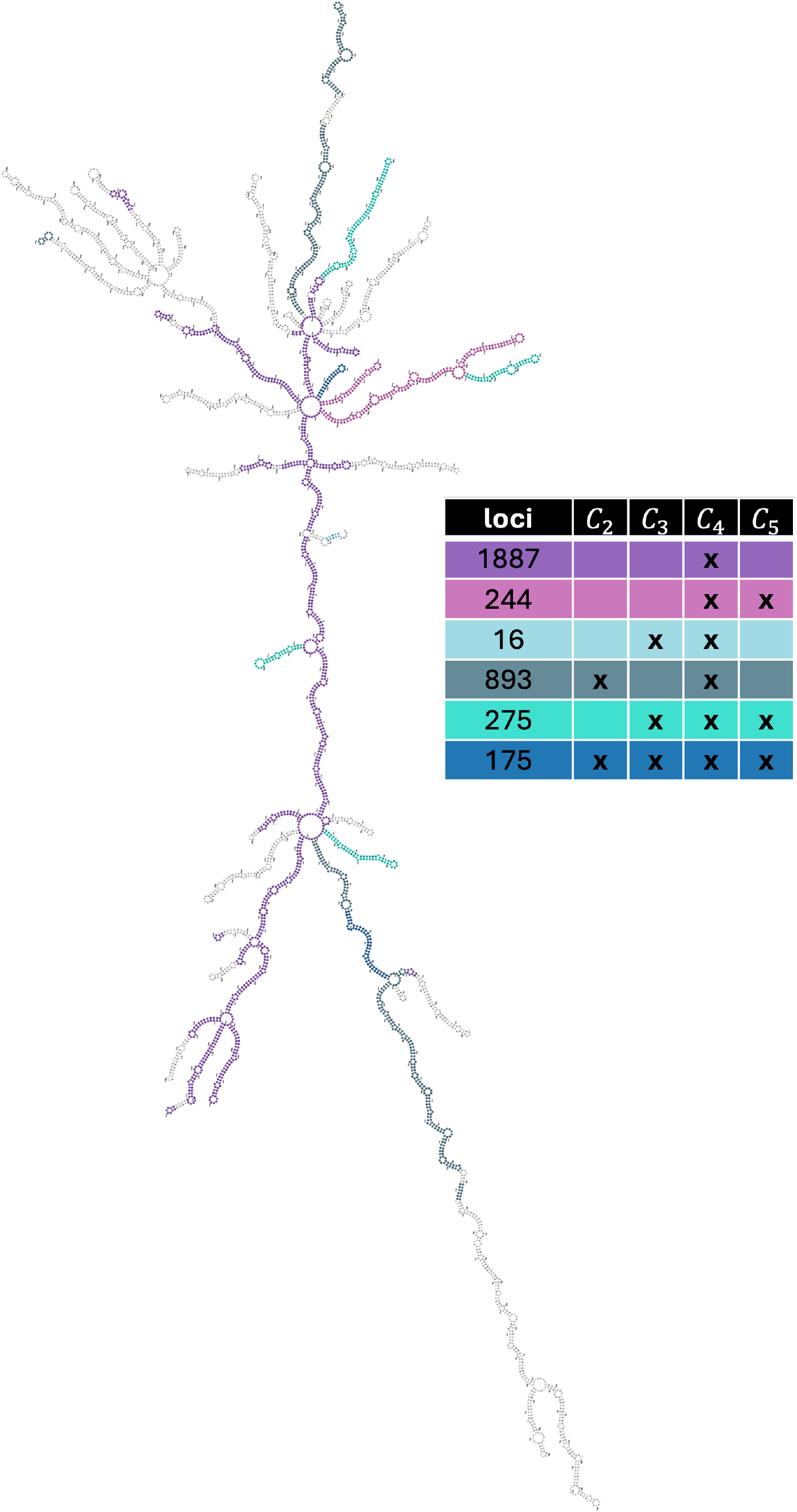
Cluster *C*_4_ of spike mRNA designs. Cluster *C*_4_ contains 2 structures, with CAI ranging from 0.799 to 0.806 and MFE from −2511.5 kcal/mol to −2500.4 kcal/mol. Table and coloring indicates cluster membership of loci with shared secondary structure.

**Figure S10:**
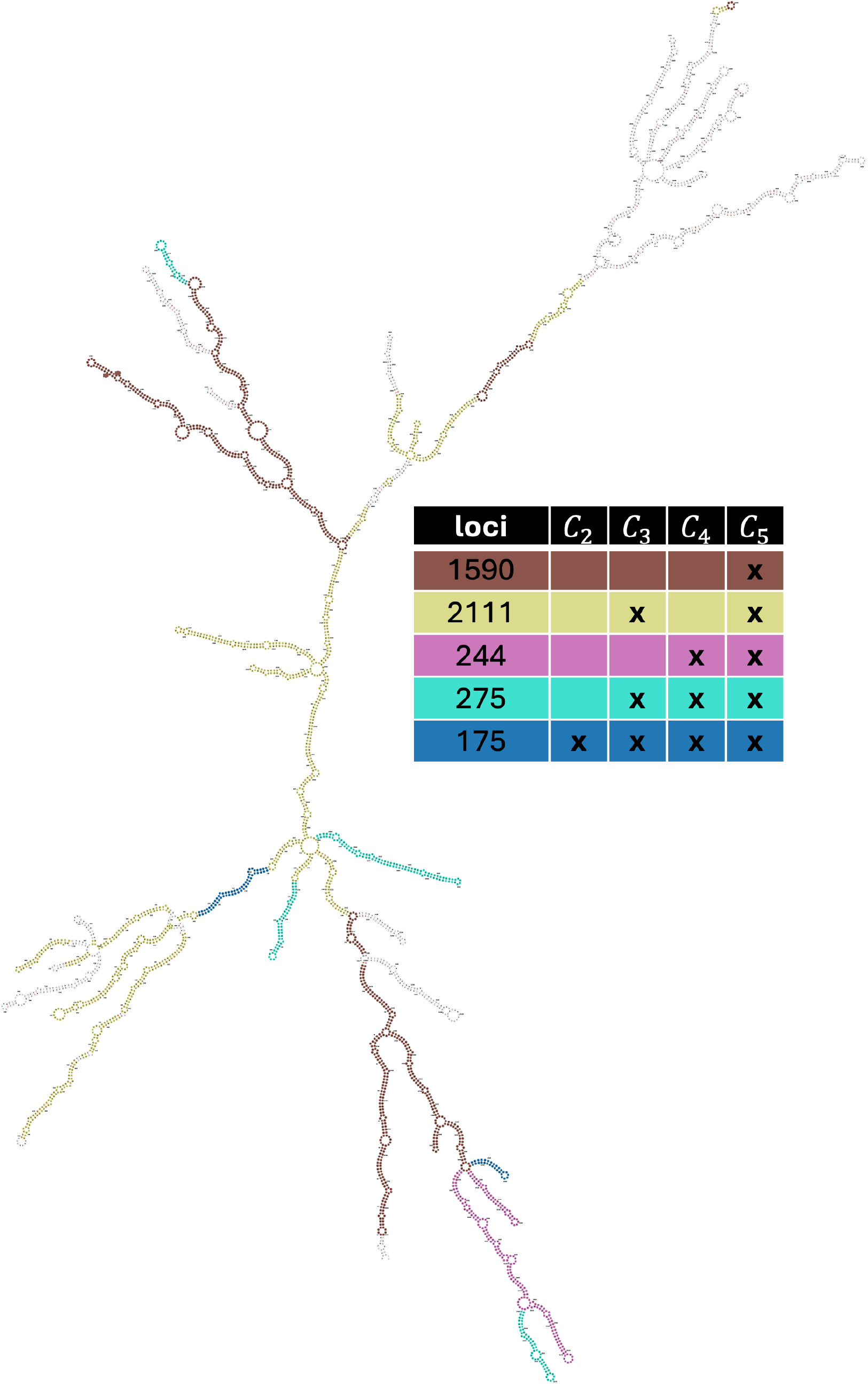
Cluster *C*_5_ of spike mRNA designs. Cluster *C*_5_ contains 16 structures, with CAI ranging from 0.717 to 0.747 and MFE from −2552.2 kcal/mol to −2548.9 kcal/mol. Table and coloring indicates cluster membership of loci with shared secondary structure.

